# Heritability of DNA methylation in threespine stickleback (*Gasterosteus aculeatus*)

**DOI:** 10.1101/2020.11.26.400531

**Authors:** Juntao Hu, Sara J. S. Wuitchik, Tegan N. Barry, Sean M. Rogers, Heather A. Jamniczky, Rowan D. H. Barrett

## Abstract

Epigenetic mechanisms underlying phenotypic change are hypothesized to contribute to population persistence and adaptation in the face of environmental change. To date, few studies have explored the heritability of intergenerationally stable methylation levels in natural populations, and little is known about the relative contribution of *cis*- and *trans-regulatory* changes to methylation variation. Here, we explore the heritability of DNA methylation, and conduct methylation quantitative trait loci (meQTL) analysis to investigate the genetic architecture underlying methylation variation between marine and freshwater ecotypes of threespine stickleback *(Gasterosteus aculeatus).* We quantitatively measured genome-wide DNA methylation in fin tissue using reduced representation bisulfite sequencing of F1 and F2 crosses, and their marine and freshwater source populations. We identified cytosines (CpG sites) that exhibited stable methylation levels across generations. We found that genetic variance explained an average of 24 to 35% of the methylation variance, with a number of CpG sites possibly autonomous from genetic control. Finally, we detected both *cis*- and *trans-*meQTLs, with only *trans*-meQTLs overlapping with previously identified genomic regions of high differentiation between marine and freshwater ecotypes, as well as identified the genetic architecture underlying two key CpG sites that were differentially methylated between ecotypes. These findings demonstrate a potential role for DNA methylation in facilitating adaptation to divergent environments and improve our understanding of the heritable basis of population epigenomic variation.

## Introduction

DNA methylation is a chemical modification to DNA that typically occurs at cytosines within CpG dinucleotides in animals (Suzuki & Bird 2008). It has been suggested that DNA methylation can play a number of biological roles, including gene expression regulation (expression, repression, alternative splicing, and spurious transcription prevention), cell-fate decision, and phenotypic evolution and adaptation to divergent environments (Bird 2007; Bossdorf *et al.* 2008; Maunakea *et al.* 2010; Feil & Fraga 2012; Jones 2012; Verhoeven *et al.* 2016; Neri *et al.* 2017; Richards *et al.* 2017). Recent genome-wide studies have revealed that DNA methylation variation is widely observed between closely related animal species and populations that have adapted to ecologically divergent environments (Massicotte *et al.* 2011; Liebl *et al.* 2013; Smith *et al.* 2015; Lea *et al.* 2016; Artemov *et al.* 2017; Le Luyer *et al.* 2017; Hu *et al.* 2018; Hu *et al.* 2019; Laporte *et al.* 2019; Heckwolf *et al.* 2020). In addition, methylation variation has been shown to have a substantial heritable component that selection can act on (Lim & Brunet 2013; Heard & Martienssen 2014; Taudt *et al.* 2016). Modification of the methylome may therefore be an important mechanism underlying phenotypic variation, adaptive evolution, and possibly ecological speciation (Jaenisch & Bird 2003; Turck & Coupland 2014; Verhoeven *et al.* 2016).

While theoretical studies have suggested that the evolutionary relevance of methylation variation is partially related to its heritability, experimental studies investigating heritable DNA methylation and its role in adaptive evolution are in their initial stages (Verhoeven *et al.* 2016; Hu & Barrett 2017; Richards *et al.* 2017). Although it is clear that DNA methylation levels can sometimes be intergenerationally stable (Jablonka & Raz 2009; Daxinger & Whitelaw 2012; Heard & Martienssen 2014), results have mainly come from plant studies, and the small number of animal studies have typically used isogenic lab lines (Morgan *et al.* 1999; Rakyan *et al.* 2003; but see Nätt *et al.* 2012; Weyrich *et al.* 2016; Weyrich *et al.* 2018; Heckwolf *et al.* 2020). The homogenous genetic backgrounds of these isogenic lines may mean that they are not representative of the methylation patterns occurring in more genetically heterogeneous populations (Herman *et al.* 2014; Verhoeven & Preite 2014). In addition, most studies in non-model species have so far been limited to describing broad patterns based on anonymous markers of DNA methylation (Schrey *et al.* 2013; Hu & Barrett 2017; Richards *et al.* 2017), which has hindered understanding of the functional relevance and genetic basis of stable methylation in these species.

Methylation variation is mainly under genetic control, which can be caused by DNA sequence variation in both *cis*- and *trans*-regulatory elements (Taudt *et al.* 2016; Hu & Barrett 2017). Recently, methylation quantitative trait loci (meQTL) analysis has found both *cis*- and *trans*-acting genetic variants underlying methylation variation (Dubin *et al.* 2015; Orozco *et al.* 2015; Kawakatsu *et al.* 2016; Meng *et al.* 2016; Taudt *et al.* 2016). *Cis*-regulatory genetic variation typically affects methylation patterns of only one or a few nearby sites and is less pleiotropic, whereas genetic variants in *trans*-regulatory elements can simultaneously change the methylation levels of multiple sites (Taudt *et al.* 2016; Do *et al.* 2017; Schulz *et al.* 2017; Hannon *et al.* 2018; Gupta *et al.* 2019). However, with the exception of a few studies (Fan *et al.* 2019) almost all meQTL studies have been conducted in model species, and thus, the prevalence of *cis*- and/or *trans*-meQTLs, and their role in adaptive evolution in natural populations remains unclear.

To explore the stability of epigenetic modification between generations, and to study the genetic architecture of methylation variation between natural populations adapted to distinct environments, we used threespine stickleback (*Gasterosteus aculeatus*), an abundant fish species in both marine and freshwater habitats in the Northern Hemisphere. Since the end of the last ice age, marine stickleback colonized freshwater lake and stream habitats that were uplifted and landlocked, resulting in replicate freshwater populations that show repeated evolution of a suite of locally adapted traits (Bell & Foster 1994). The repeated adaptive divergence between marine and freshwater populations makes this a powerful system to study the ecology and genetic architecture of adaptation (Jones *et al.* 2012). In the last decade, a variety of genetic and genomic resources have been developed for this species (Baird *et al.* 2008; Hohenlohe *et al.* 2010; Jones *et al.* 2012; Ishikawa *et al.* 2017; Peichel & Marques 2017). In addition, genome-wide methylation variation between marine and freshwater populations (Smith *et al.* 2015) and between males and females (Metzger & Schulte 2018) have been characterised, as well as the demonstration of methylation responses to environmental change (Artemov *et al.* 2017; Metzger & Schulte 2017; Heckwolf *et al.* 2020). However, the intergenerational stability of methylation in stickleback, and the genetic architecture underlying methylation variation between marine and freshwater ecotypes, remain unclear.

We address these gaps by performing an epigenomic survey of fin tissue from sticklebacks under a common garden experimental design with controlled crosses. We first examined methylation divergence between marine and freshwater ecotypes. We then explored levels of methylation and its genetic basis across two generations of the marine-freshwater hybrid lines, and performed meQTL analysis with two F2 families to characterise the genetic architecture of methylation variation between ecotypes. We investigate four specific questions: (1) Is variation in DNA methylation stable between generations? (2) What is the genetic heritability of intergenerationally stable CpG sites? (3) What is the genetic architecture of DNA methylation differences between the stickleback ecotypes? (4) What are the relative contributions of *cis*- and *trans*-meQTLs to DNA methylation differences? Answering these questions will help to provide a baseline for understanding the heritability of methylation variation, and the role of methylation variation in facilitating population persistence and potentially local adaptation in natural populations.

## Materials and methods

### Sampling and husbandry

We collected adult threespine stickleback from one marine (Bamfield Inlet, BI, 48°49’12.69”N, 125° 8’57.90”W), and two freshwater (Hotel Lake, HL, 49°38’26.94”N, 124° 3’0.69”W, and Klein Lake, KL, 49°43’32.47”N, 123°58’7.83”W) locations in Southwestern British Columbia, Canada in May 2015 (Fig. 1). We transported all fish to an aquatic facility at the University of Calgary, and separated them into population-specific 113 L glass aquaria. We maintained a common garden environment at a density of approximately 20 fish per aquarium, salinity of 4-6 ppt, water temperature of 15 ± 2 °C, and a photoperiod of 16 L: 8 D for one year before making crosses. This period of time should minimize any effects of transportation and allow sufficient time for marine populations to acclimate to hypoosmotic conditions (McCairns & Bernatchez 2010; Morris *et al.* 2014; Wang *et al.* 2014; Artemov *et al.* 2017). We kept each aquarium as a closed system with its own filter, air pump, water supply, and temperature regulator. We fed all fish *ad libitum* once per day with thawed bloodworms (Hikari Bio-Pure Frozen Bloodworms).

**Fig. 1.**
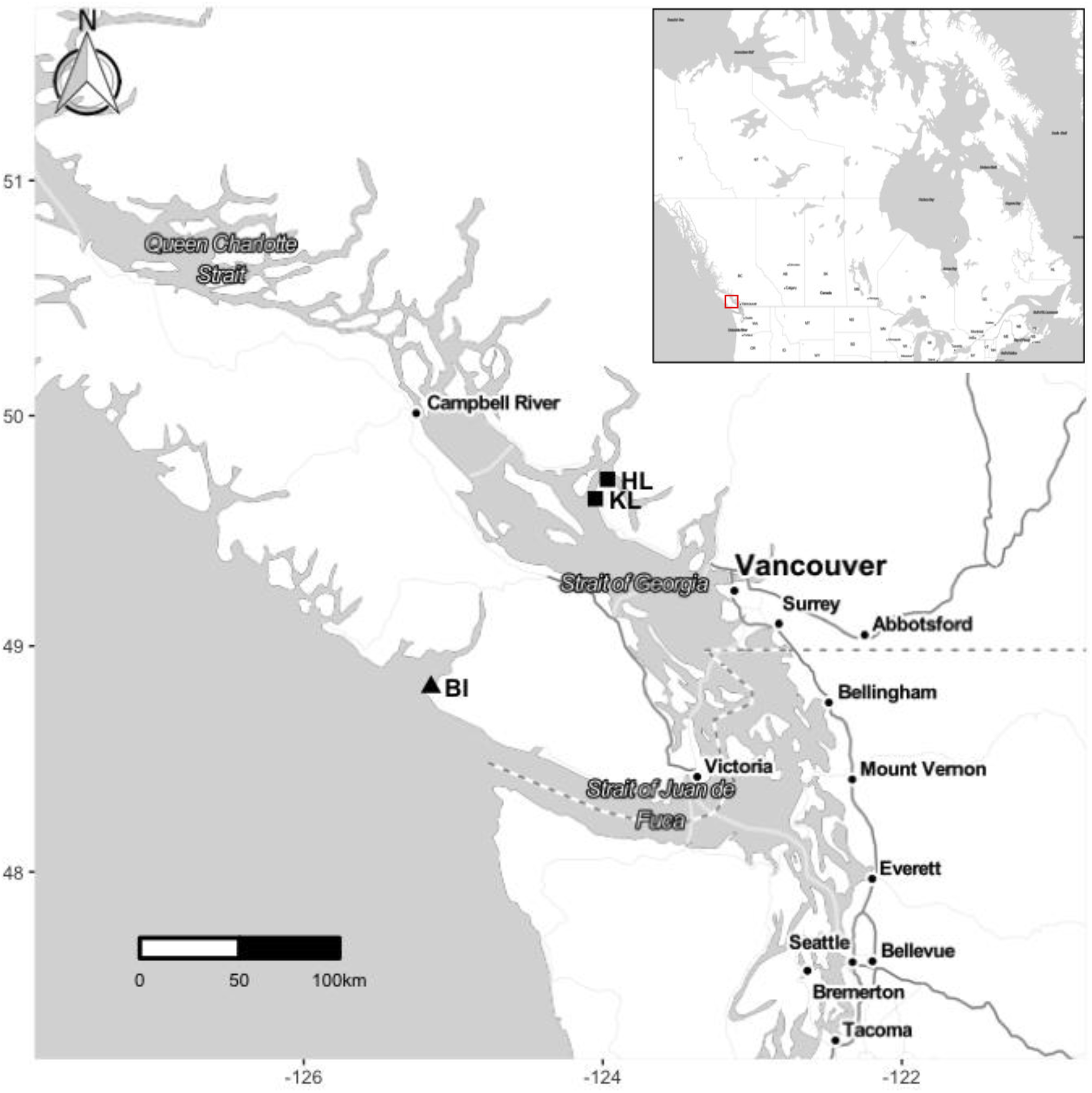
Geographical location of threespine stickleback populations used in this experiment. Triangle indicates the marine sampling site, squares indicate freshwater sampling sites. BI, Bamfield Inlet (marine); HL, Hotel Lake (freshwater); KL, Klein Lake (freshwater). The red square in the inset shows the location of sampling sites in relation to the broader geographic region (the west coast of Canada).

### Crossing design

Threespine stickleback are typically found in either marine or freshwater habitats, but distinct marine and freshwater ecotypes can hybridize, which can facilitate the detection of associations between genotype and phenotype (Jones *et al.* 2012). We generated genetically heterogeneous marine-freshwater F1 families from wild-caught parents by collecting eggs from one marine female and extracting testes from one freshwater male per cross (Fig. S1). To generate a cross, we first equally distributed the eggs into a Petri dish containing fresh water. We then euthanized the male using an overdose of eugenol and removed the testes. We crushed the testes in a Petri dish, with the water activating the released sperm and allowing fertilization. Fertilized eggs were left within the Petri dish for 20 minutes before being suspended in a well-aerated mesh-bottom container within 37 L glass aquaria, with an air stone for oxygenation and a sponge filter. In total, we produced one F1 family of BIxHL hybrids (hereafter referred to as HL_F1), and three F1 families of BIxKL hybrids (hereafter referred to as KL_F1). After hatching, the larval fish from the same family were reared in the same 37 L aquaria until reaching approximately 1 cm total length (TL), at which time the families were equally split into aquaria to maintain low densities. The fish and fry were fed twice daily with live *Artemia spp.* nauplii. At approximately 2 cm TL, juvenile stickleback fish were transitioned to a diet of chopped thawed bloodworms once per day *ad libitum.* They were then transitioned to an adult diet of full thawed bloodworms gradually. We sampled caudal fin clips (hereafter referred to as ‘fin clips’) when individuals reached a 3.5 cm TL or more. In addition to the fish we used to generate the F1 crosses, we also sampled extra parental fish from the same marine or freshwater population. Fin clips were stored in 70% ethanol in microcentrifuge tubes at room temperature until extraction of genetic material.

To generate F2 families, we randomly selected and crossed one male and one female sibling within an F1 family from each hybrid line (HL or KL) using the same crossing methods. We produced one F2 family of HL hybrids (hereafter referred to as HL_F2) and one F2 family of KL hybrids (hereafter referred to as KL_F2). Fish were raised as described above. We randomly selected fish from HL_F2 and KL_F2 families, and sampled fin clips when individual reached approximately 3.5 cm TL. We stored all fin clips as described above. In addition to the fish we used to make the F2 crosses, we also randomly sampled extra F1 fish from all F1 families. In total, we sampled 94 fish, including 11 parental fish (six marine females; two HL and three KL freshwater males), 19 F1 fish (7 HL_F1 and 12 KL_F1), and 64 F2 fish (28 HL_F2 and 36 KL_F2). Detailed information about sex and family is included in Table S1. All sampling, crossing, and housing protocols were approved by the University of Calgary Life and Environmental Science Animal Care Committee (AC13-0040 and AC17-0050) following the ethical standards maintained by the Canadian Council for Animal Care.

### Tissue choice

The choice of tissue used for genome-wide mapping of cytosines can influence the interpretation of methylation patterns (Stricker *et al.* 2017). We conducted our analyses using caudal fin tissue for several reasons. It has been shown that fin position, caudal depth, caudal fin size are different between marine and freshwater stickleback, and that this phenotypic difference is heritable and associated with repeated adaptation to divergent marine and freshwater environments (Walker 1997; Jones *et al.* 2012). Because methylation is tissue-specific, choosing a tissue showing phenotypic differences between ecotypes increases the likelihood of finding meQTLs that contribute to this ecotype divergence. Caudal fins can also be dissected quickly and consistently, and the excision of fin tissue does not affect survival.

### DNA extraction and sex determination

We extracted DNA from caudal fin using phenol:chloroform:isoamyl alcohol (25:24:1), and assessed the quality and quantity using Tecan Infinite^^®^^ 200 NanoQuant and Quant-iT PicoGreen^®^ dsDNA assay kit (ThermoFisher Scientific). We determined the sex of fish following Peichel *et al.* (2004).

### Reduced representation bisulfite sequencing

To measure genome-wide DNA methylation levels, we used reduced representation bisulfite sequencing (RRBS) (Meissner *et al.* 2008; Gu *et al.* 2011), following Boyle *et al.* (2012) with some minor modifications. For each individual, we created a library from 120 ng of genomic DNA, and ligated the *MspI*-digested fragments in each library with unique Illumina TruSeq adapters. We targeted fragments of 160-340bp (including ~120bp of adapter sequence) using NaCl-PEG diluted SpeedBeads (Rohland & Reich 2012). We split the libraries into four pools (three pools of 24 libraries and one pool of 22 libraries), and treated the pools with sodium bisulfite (EpiTect, Qiagen) following a protocol for formalin-fixed paraffin-embedded samples (Gu *et al.* 2011). After two rounds of bisulfite treatment to ensure complete conversion of unmethylated cytosines, each pool was amplified with Illumina primers, and loaded in four lanes (100-bp single-end reads) of a Hiseq 2500 at the McGill University and Genome Quebec Innovation Centre. In total, we sequenced all 94 fish sampled across three generations (Table S1). Each sample was sequenced to a mean depth (± SD) of 8.094 ± 2.532 million reads. The average mapping efficiency was 61.4 ± 4.7% (± SD). We quantified methylation at non-CpG motifs and found less than 1% non-CpG cytosines were methylated, suggesting a highly efficient bisulfite conversion.

### Read filtering and mapping

To remove adapter contamination, low-quality bases, and bases artificially introduced during library construction, we trimmed reads using Trim Galore! v0.4.4 (https://www.bioinformatics.babraham.ac.uk/projects/trim_galore/), with the ‘rrbs’option. We then used the program Bowtie2 v2.2.9 (Langmead & Salzberg 2012), implemented in Bismark v0.17.0 (Krueger & Andrews 2011) to align trimmed reads for each sample to the stickleback genome (ENSEMBL version 98) with default settings, except for tolerating one non-bisulfite mismatch per read. We only included reads that mapped uniquely to the reference genome, and cytosines that had at least 10x coverage in downstream analyses. Only CpG context cytosine methylation was analysed because CpG methylation is the most common functional methylation in vertebrates (Suzuki & Bird 2008).

### General methylation patterns

To identify general methylation patterns, we first performed a principal component analysis (PCA) on methylation levels in all samples using the *prcomp* function in R (R Core Team, 2018, v3.4.3). We ran the analysis by first identifying cytosines that were covered in all samples using the R package methylKit v1.4.1 (Akalin *et al.* 2012). Read coverage was then normalized between samples, using the median read coverage as the scaling factor. A minimum of ten reads in all samples was required at a CpG site for that site to be analysed. We removed CpG sites that were in the 99.9^th^ percentile of coverage from the analysis to account for potential PCR bias. We calculated the methylation levels by extracting the total amount of methylation-supporting reads, and the total coverage of each CpG site, using the *percMethylation* function in the R package methylKit. To improve methylation estimates, we corrected for SNPs, which could have resulted in an incorrect methylation call if C-to-T and G-to-A SNPs were falsely interpreted as unmethylated cytosines, following Heckwolf *et al.* (2020). We first identified SNPs using the methylation value of each CpG site of all 11 parental individuals for input to Bis-SNP v0.82.2 (Liu *et al.* 2012) with the default parameters. Because Bis-SNP is sensitive to the directionality of the RRBS protocol (i.e., whether sequenced reads come from the original forward and reverse strands when calling C-to-T and G-to-A SNPs), we used a directional bisulfite-seq protocol that is similar to Krueger *et al.* (2012). We observed a similar number of reads per individual in our study vs. Heckwolf *et al.* (2020), suggesting that sufficient coverage on both strands was obtained to distinguish SNPs from conversions at C-to-T and G-to-A SNPs. We chose parental samples for identifying SNPs because they are the genetic source of the F1s and F2s, and are the most genetically heterogeneous samples. We used GATK’s VariantFiltration and SelectVariants to restrict variants to diallelic sites, and filter variants based on the following GATK variant annotation cut-offs: QD < 2.0, MQ < 40.0, MQRankSum < −12.5, and ReadPosRankSum < −8.0. We then used VCFtools v0.1.16 (Danecek *et al.* 2011) to remove SNPs with a minor allele frequency (MAF) greater than 0.0049 (Heckwolf *et al.* 2020), and more than 10% missing data across all 11 parental samples. Using MAF thresholds from 0.001 to 0.01 resulted in similar numbers of filtered SNPs. We calculated pairwise weighted *Fst* between BI, HL and KL fish in the parental generation using VCFtools on filtered SNPs. We produced a list of positions (C-to-T and G-to-A SNPs) for correcting methylation estimates, using custom written Perl scripts from Heckwolf *et al.* (2020) and the R package GenomicRanges v.1.30.3 (Lawrence *et al.* 2013). In addition, it has been suggested that sex specific methylation affects less than 0.1% of CpG sites on autosomal chromosomes, but more than 5% of CpGs on the sex chromosome in stickleback (Metzger & Schulte 2018). Therefore, to exclude a potential sex bias, we removed all CpGs located on the sex chromosome (group XIX). In total, we retained 52,940 CpG sites that passed the filtering step. To perform PCA, methylation levels at each CpG site were taken as input variables, whereas each point in multidimensional space represented a stickleback individual. Finally, to compare DNA methylation variation levels between F1 and F2 fish in each hybrid line, we calculated DNA methylation levels in 7,840 1-kb tiling windows (step = 1 kb; size = 1 kb) compiled from the same 52,940 CpG sites, and compared the standard deviations of methylation levels for each genomic window within each hybrid line.

### Analysis of methylation divergence between ecotypes

To examine methylation divergence between ecotypes, we performed a differential methylation analysis between marine and freshwater populations from the parental generation, using the 52,940 CpG sites that passed the filtering step above. CpG sites were considered to be differentially methylated cytosines (DMCs) with a false discovery rate correction *Q*-value < 0.01 and a minimum required methylation difference of 15% between ecotypes, using the R package methylKit with sampling site (i.e., BI for the marine population, and KL and HL for the freshwater populations) as an covariate. We visualized differential methylation patterns across individuals and obtained clustering of samples and DMCs in heatmaps with the “complete” clustering method on Euclidian distances, using the R package pheatmap version 1.0.8 (https://cran.r-project.org/web/packages/pheatmap/index.html). We clustered hyper- and hypomethylated DMCs between ecotypes using relative percent DNA methylation, which is the normalized percent DNA methylation scaled for each DMC’s percent DNA methylation (median percent methylation as 0) of marine and freshwater fish in heatmaps. We also clustered individual fish based on overall methylation patterns across DMCs. We then analysed the proportion of cytosines within genomic features (promoter/exon/intron/intergenic; promoters are defined as regions being upstream 1000 bp and downstream 1000 bp around the transcription start sites (TSSs)) for DMCs, using the R package genomation v1.6.0 (Akalin *et al.* 2015). Because *MspI* restriction sites are not randomly distributed in the genome, we built a null distribution of genomic features based on all filtered CpG sites (i.e., 52,940 CpG sites). We gave precedence to promoters > exons > introns > intergenic regions when features overlapped (Smith *et al.* 2015; Hu *et al.* 2018). Finally, we annotated genes associated with DMCs, using the R packages biomaRt v2.34.2 (Durinck *et al.* 2005; Durinck *et al.* 2009) and ChIPpeakAnno v3.12.7 (Zhu *et al.* 2010; Zhu 2013) on the stickleback reference genome from Ensembl 98 database, and performed gene ontology (GO) analysis on DMC-associated genes, using the R package topGO v2.28.0 (Alexa *et al.* 2006). Over-represented GO terms were those with multiple-test corrected *P*-values (Benjamini-Hochberg’s false discovery rate) below 0.1, based on a Fisher’s exact test. We compared DMC-associated genes with the genes associated with the 52,940 CpGs that passed the filtering step.

### Analysis of intergenerationally stable methylation

We considered a CpG as intergenerationally stable when the CpG was not significantly differentially methylated between F1 and F2 generations of the same family within the same hybrid line (HL or KL) and fulfilled this criterion in both hybrid lines. Note that these ‘stable’ sites are not necessarily ‘heritable’ in the sense of methylation variation between individuals being due to additive genetic factors. DMCs between fish in F1 vs. F2 generations were identified using the same method as described above for identifying DMCs between ecotypes, with sequencing lane as a covariate. We identified 137 and 82 DMCs within HL and KL hybrid lines, respectively. These sites were removed from the 52,940 CpG sites that passed the filtering step to provide the dataset of intergenerationally stable sites. We clustered fish based on the similarity of their DNA methylation profiles, with the “ward” clustering method on Pearson’s correlation distances, using the *clusterSamples* function in the R package methylKit. We also compared the locations of DMCs between ecotypes with the locations of intergenerationally stable CpG sites to assess which of the sites involved in methylation divergence between ecotypes are stable across generations.

Our criterion for identifying CpG sites with intergenerationally stable methylation is such that a type 2 error in our differential methylation test between generations (a false negative for differential methylation between F1 and F2 within a line) will lead to a false positive for stable methylation. To investigate the potential importance of this type of error in our data, we conducted a power analysis using a simulated data set in methylKit, following Wreczycka *et al.* (2017) with some minor modifications. To be conservative, our simulated dataset consisted of eight samples (four F1s and four F2s, matching the minimum number of fish in a line from each generation in the empirical dataset). We modelled the read coverage following a binomial distribution and defined the methylation levels following a beta distribution with parameters alpha = 0.4, beta= 0.5 and theta =10. We ran simulations of differential methylation at 1% of 52,940 CpG sites, with effect sizes of 5%, 10%, 15%, 20% and 25% differential methylation, respectively. After correcting for the covariate of sequencing lane (the same sequencing lane was assigned to two samples within F1 or F2 generation, for a total of four sequencing lanes), we adjusted *P*-values for multiple testing using the *Q*-value method (Storey & Tibshirani 2003), and considered CpGs to be DMCs with a false discovery rate correction *Q*-value < 0.01. Finally, we calculated the proportion of CpGs that were falsely identified as non-DMCs (false negatives) among all CpG sites under each effect size (5% to 25%) above.

We distinguished between three categories of methylated sites that were stable between F1 and F2 generations: 1) constitutively hypermethylated sites, which are CpG sites with average DNA methylation levels greater than 0.9 in all samples, 2) constitutively hypomethylated sites, which are CpG sites with average DNA methylation levels less than 0.1 in all samples (Lam *et al.* 2012; Lea *et al.* 2016), and 3) methylated sites with average DNA methylation levels between 0.1 and 0.9 (hereafter referred to ‘variable sites’). Finally, we analysed the proportion of cytosines within genomic features for CpG sites in each category and annotated genes associated with all intergenerationally stable CpG sites by testing for overlap between the locations of CpG sites and genomic regions of genes, following the same method as described above for ecotype DMCs.

### Heritability of stable methylation

To determine the genetic heritability of the intergenerationally stable CpG sites across F1 and F2 generations, we first identified SNPs using the aligned reads of all F1 and F2 individuals for input to Bis-SNP, following the same SNP calling and filtering steps as described above with some minor modifications. We retained three sets of SNPs by filtering the 92,983 SNPs using a constant MAF cut-off (0.005) and three missing data cut-offs (10%, 30%, 50%) across all F1 and F2 individuals. We used BCFtools (https://github.com/samtools/bcftools) to exclude sites that were under linkage disequilibrium (LD, pairwise r^2^ > 0.8 within a window of 1Mb) or on the sex chromosome. Finally, we used a linear mixed model implemented in PyLMM (http://genetics.cs.ucla.edu/pylmm/index.html) to test whether variation at SNPs is significantly associated with methylation levels at stable CpG sites in F1 and F2 generations, after correcting for sequencing lane variation and kinship based on the SNP data. We adjusted multiple-test *P*-values using Benjamini-Hochberg’s false discovery rate and considered an association to be significant when the corrected *P*-value < 0.05.

We estimated the narrow sense heritability of DNA methylation levels for individual CpG sites of all F1 and F2 individuals using a linear mixed model approach (Yang *et al.* 2010) implemented in the R package lmmlite (https://github.com/kbroman/lmmlite). We treated the methylation levels at individual CpG sites of all F1 and F2 individuals as phenotypes, and assumed each phenotype **y** can be modelled as **y = 1_n_μ + u + e**, where the random variable **u** follows a normal distribution centred at zero with variance σ_g_^2^K, and **e** represents an independent noise component with variance σ_e_^2^. The matrix K is the same kinship matrix as calculated above. For each trait we estimated σ_g_^2^ and σ_e_^2^ using the restricted maximum likelihood (REML) approach, with correction for the covariate of sequencing lane, and calculated the heritability as h^2^ = σ_g_^2^/(σ_g_^2^ + σ_e_^2^). Finally, we calculated the average heritability by taking the mean of heritability values of all CpG sites.

### meQTL analysis

To identify the genetic architecture of methylation divergence between marine and freshwater stickleback, we performed meQTL mapping of the methylome in two F2 families of marine-freshwater hybrids. Due to the distinct methylation patterns that may be caused by genetic variation between the two hybrid lines (Fig. S2), we performed mapping separately for each hybrid line. We first filtered SNPs that were not located on the sex chromosome, had less than 10% missing data, and had low LD in HL_F2 or KL_F2 samples, using the same SNP filtering steps as described above. We then compiled a percentage methylation level matrix among HL_F2 or KL_F2 samples containing the 52,940 CpG sites that passed these filtering steps.

Finally, we truncated these sites by the 10% minimum range of methylation variation across samples to reduce non-informative sites that could possibly inflate test statistics and create spurious SNP-CpG pairs. After filtering, we retained 525 SNPs and 27,614 CpGs in HL_F2, and 330 SNPs and 27,039 CpGs in KL_F2 for meQTL analysis, with no overlap between the retained SNPs in each line. We tested all genome-wide SNP-CpG pairs using the R package MatrixEQTL v2.2 (Shabalin 2012). This package enables rapid computation of QTLs by only retaining those that are significant at a pre-defined threshold. We fit an additive linear model to test if the number of alleles (coded as 0, 1, 2) predicted percentage DNA methylation levels (value ranging from 0 to 1) at each CpG site, including sequencing lane as a covariate. We used a Bonferroni-corrected multiple-test corrected threshold, set it to genome-wide significance for GWAS and divided by the number of CpG sites tested (i.e., HL_F2: 5 x 10^-8^/27,614 = 1.81 x 10^-12^; KL_F2: 5 x 10^-8^/27,039 = 1.85 x 10^-12^). We chose this stringent threshold to call meQTLs to minimize the possibility of false positives (Orozco *et al.* 2015). We calculated the distance between a SNP and a CpG site within a significant meQTL and defined a SNP as *cis*-acting if the SNP was located within 1Mb from its associated CpG site or *trans-*acting if the SNP was located more than 1Mb from its associated CpG site (Zhang *et al.* 2014). We then performed GO analysis on genes associated with unique SNPs within significant meQTLs and identified over-represented GO terms, using the same method as described above. The gene pools against which we compared the unique SNPs were the genes associated with the SNPs that passed the filtering step. Because previous studies have suggested that meQTLs and eQTLs are likely to co-occur in close genomic proximity, we compared locations of significant meQTLs in our study to significant eQTLs identified in Ishikawa *et al.* (2017), which also used a marine-freshwater hybrid design. Ishikawa *et al.* (2017) identified eQTLs under a range of salinities, but we only used eQTLs that they identified under 3.1 ppt, which is a similar salinity level to our experimental conditions. Finally, to investigate the role of meQTLs in adaptation to different habitats in stickleback, we compared locations of unique SNPs within significant meQTLs to previously documented regions of parallel genomic divergence between marine and freshwater sticklebacks (Hohenlohe *et al.* 2010; Jones *et al.* 2012; Terekhanova *et al.* 2014), and identified genetic architecture of DMCs between ecotypes by comparing the locations of DMCs and unique CpG sites associated with significant meQTLs. In addition to performing the meQTL analyses using MatrixEQTL as described above, we also validated meQTL results within each hybrid line using the R package R/qtl v.1.46-2 (Broman *et al.* 2003) with default settings, except for rescaling the basepair positions of SNPs by multiplying by a constant of 3.11 x 10^-6^ due to the genome-wide recombination rate of 3.11cM/Mb in stickleback (Roesti *et al.* 2013). We calculated genomewide logarithm of the odds (LOD) thresholds through 1,000 permutations, using the n.perm function in the R package R/qtl and set the 95^th^ percentile LOD score as the significance threshold (Hoglund *et al.* 2020).

### Data Accessibility

Raw Illumina sequencing reads for the 94 analysed individuals can be downloaded from the NCBI Short Read Archive (BioProject ID: PRJNA587332). The cytosine coverage files (.cov) for the 94 analysed individuals and codes used for analyses in this study are available through Github (https://github.com/barrettlabecoevogeno/Heritability_DNA_methylation_sticklebacks). Additional supplemental material is available at figshare.

## Results

### General methylation patterns

To identify general methylation patterns, we performed principal component analyses (PCA) on the methylation levels of filtered CpG sites represented in all samples (Fig. 2). PC1 reflected sequencing lane chemistry (Fig. S2; Table S2) and so we included sequencing lane as a covariate in all downstream analyses. When analysing all samples, PC2 (variance explained: 5.1%) clearly separated parental and F2 samples, with F1 samples filling the intermediate space between parental and F2 samples. Furthermore, the PCA separated the samples by sire (HL vs. KL) in F1 populations along PC3, which accounted for 3.2% of the variance observed in the data set (Fig. 2a). In the parental fish, the PCA separated samples mainly by their habitat along PC3 (Fig. 2b), whereas the F1 generation shows clustering based on family (Fig. 2c). The PCA also revealed some clustering between the HL and KL hybrid lines in the F2 generation, although there is no clear separation between lines (Fig. 2d). Within families, we found significantly higher mean DNA methylation variance in the F2 generation than the F1 generation of families in the HL line *(W* = 2.94 × 10^7^; *P* = 5.61× 10^-6^), but not the KL line (*W* = 3.08 × 10^7^; *P* = 0.775).

**Fig. 2.**
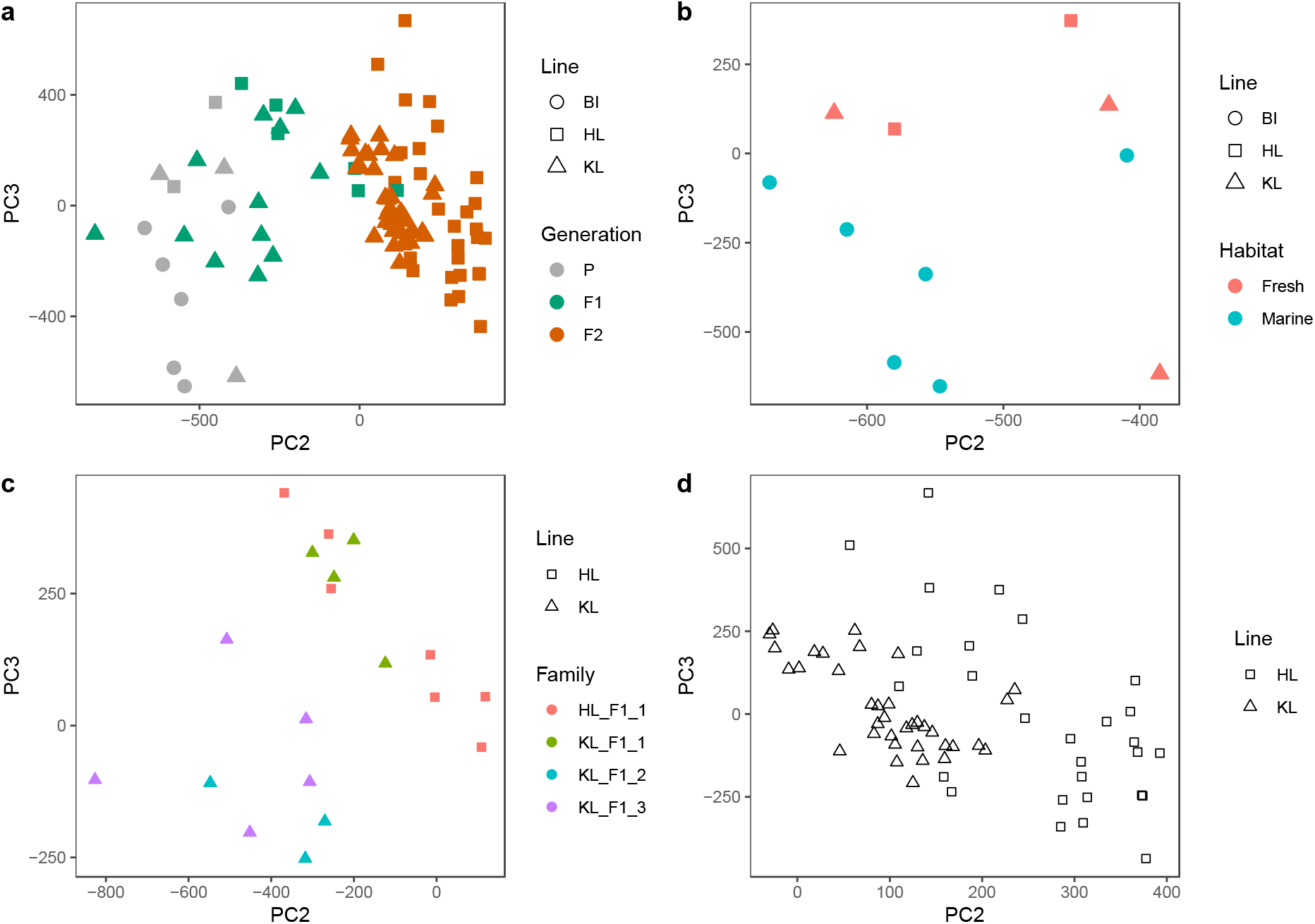
Principal component analysis (PCA) of DNA methylation profiles based on all CpG sites after filtering (See Methods) in a) all individuals from parental, F1 and F2 generation, b) parental generation, c) F1 generation, and d) F2 generation. Line: Sampling site of parental fish in generation P, parental sire of fish in the F1 generation, and grandparental sire of fish from the F2 generation.

The average pairwise *Fst* calculated between populations was 0.03 (BI vs. HL), 0.04 (BI vs. KL) and 0.01 (HL vs. KL). These values are comparable to what has been reported between other marine and freshwater populations of stickleback using SNPs extracted from RAD-seq (e.g., Hohenlohe *et al.* 2010; Catchen *et al.* 2013; Lescak *et al.* 2015; Garcia-Elfring *et al.* Unpublished) and whole genome sequencing (Shanfelter *et al.* 2019), suggesting that our use of SNPs identified from RRBS should not bias estimates of genetic differentiation relative to other methods.

### Methylation divergence between ecotypes in the parental generation

We identified 891 DMCs between parental fish sampled from marine vs. freshwater habitats after false discovery rate correction. Based on Euclidean distances, individual fish clustered by their ecotypes, with the freshwater fish further clustered by their sampling site (HL vs. KL; Fig. 3a). When analysing the methylation patterns of the 891 CpGs across generations, we found two major clusters, with the first cluster only including marine fish from the parental generation, and the second cluster including all freshwater fish from the parental generations and all F1 and F2 hybrids (Fig. S3). When comparing the mean methylation levels of the 891 ecotype DMCs in F1s versus parents, we found that in both hybrid F1 lines, there was a significantly greater number of CpG sites with methylation levels that were intermediate between the values of the parents than the number of CpG sites with methylation levels outside the values observed in the parents (KL_F1: *G* = 294, df = 1, *P* < 2.20 × 10^-16^; HL_F1: *G* = 124, df = 1, *P* < 2.20 × 10^16^). In the F2 generation, we observed a greater proportion of sites showing methylation values outside the range observed in their F1 parents relative to the pattern between F1s and their wild parents (KL_F2: *G* = 241, df = 1, *P* < 2.20 × 10^-16^; HL_F2: *G* = 9.72, df = 1, *P* = 1.82 × 10^-3^). In addition, we found a marginally greater proportion of sites showing bias in methylation levels towards those of the mother in HL_F1 (*G* = 4.10, df = 1, *P* = 0.0428) but not KL_F1 (*G* = 0.576, df = 1, *P* = 0.448) when compared to the parental generation. However, sex is confounded with parental habitat in this comparison (marine fish are always female, and freshwater fish are always male). This confounding effect is not present in the F1 generation, where the same analysis found a marginally greater proportion of sites showing bias toward the F1 mother in KL_F2 (*G* = 3.85, df = 1, *P* = 0.0497) but not HL_F2 (*G* = 0.0627, df = 1, *P* = 0.802).

**Fig. 3.**
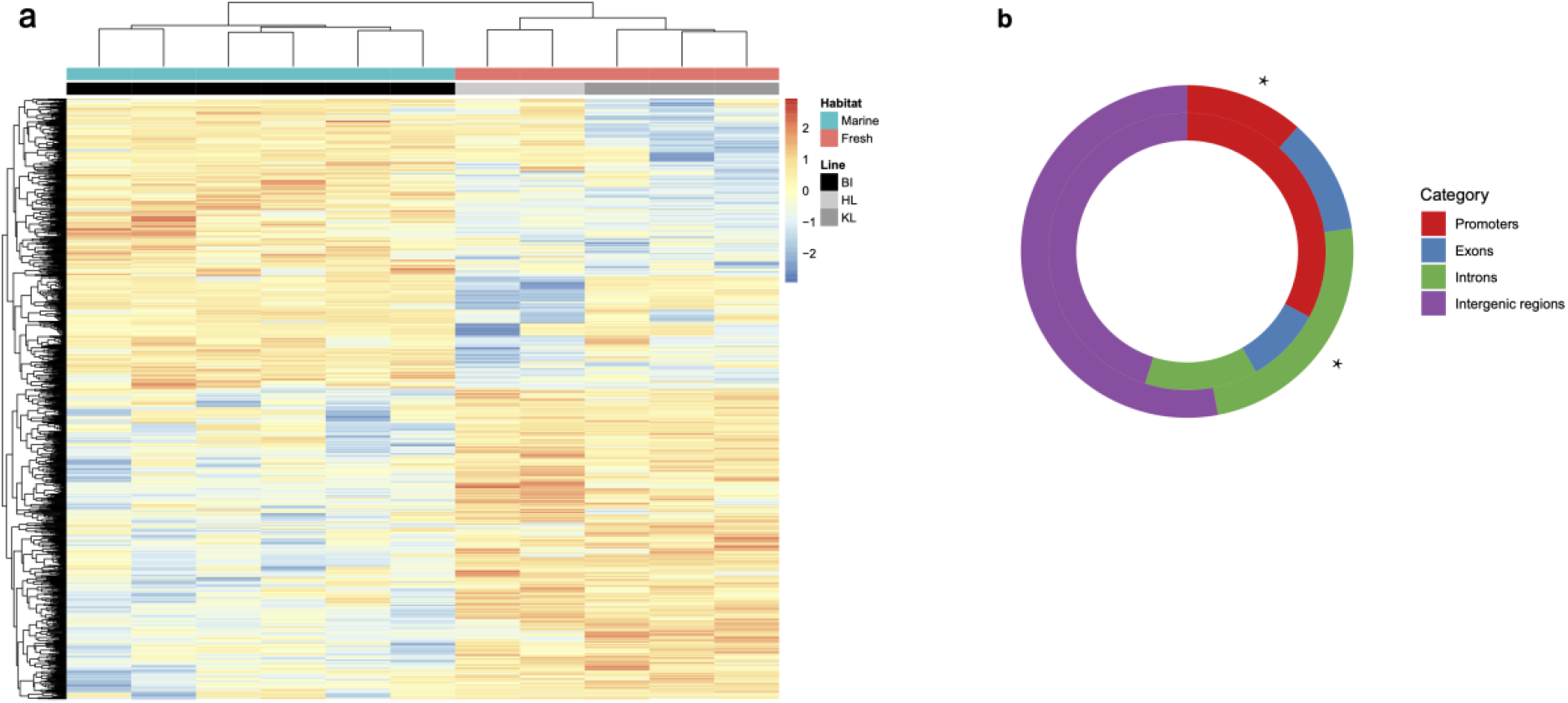
(a) Heatmap of methylation levels of the 891 DMCs between marine and freshwater ecotypes from the parental generation. Each column represents a colour-coded individual: blue for marine fish, red for freshwater fish; black for marine fish from BI, light grey for freshwater fish from HL, and dark grey for freshwater fish from KL. Each row represents one of the DMCs, which are clustered based on the similarities of the methylation patterns between individuals. Darker red indicates greater methylation in an individual for that DMC. Darker blue indicates lower methylation in an individual for that DMC. Individual dendrogram positions are based on overall methylation patterns across the 891 DMCs. (b) The proportion of genomic features (promoters, exons, introns or intergenic regions) in the 891 DMCs. Outer rings describe the locations of DMCs in each category; inner rings describe the features of null distribution of all filtered CpGs. Asterisks denote significant differences between the features of DMCs in each category vs. the features of null distribution of CpGs across the genome using a *G* test at *P* < 0.01.

Identified DMCs between ecotypes showed no significant bias towards hyper-versus hypomethylation (430 hypermethylated and 461 hypomethylated DMCs; *G* = 1.21 × 10^-3^, df = 1, *P* = 0.972). However, these DMCs showed significant enrichment within introns when compared to the null distribution of all filtered sites (introns: *G* = 8.87, df = 1, *P* = 2.90 × 10^-3^; Fig. 3b). In addition, we found no overlap between the locations of the 891 DMCs and the locations of sex-biased DMCs identified in Metzger & Schulte (2018), suggesting that removing CpGs located on the sex chromosome effectively minimized any potential sex-biased differential methylation. In total, DMCs mapped to 228 genes, with some genes having been shown to be associated with differential expression or methylation between ecotypes in recent studies (e.g., differentially expressed genes in gill: *atp1a2a, g6pd,* Artemov *et al.* 2017; differentially methylated genes in fillet: *g6pd, chchd3a,* Smith *et al.* 2015). GO analysis showed no significant GO term enrichment.

### Intergenerationally stable methylation

We found that 99.6% of CpG sites (52,729 out of 52,940) were not differentially methylated across generations in both lines, suggesting the vast majority of sites show stable levels of methylation across generations. Our power analysis suggests that a small proportion of sites (less than 1%) are likely to have been falsely identified as non-DMCs (Fig. S4) across all effect size groups, suggesting the influence of type 2 error on our criterion for calling stable methylation would only effect a small number of sites. Based on Pearson’s correlation distance calculated from the 52,729 CpG sites, most individuals clustered by generation (F1 vs. F2) and by hybrid line (HL vs. KL) (Fig. 4a).

**Fig. 4.**
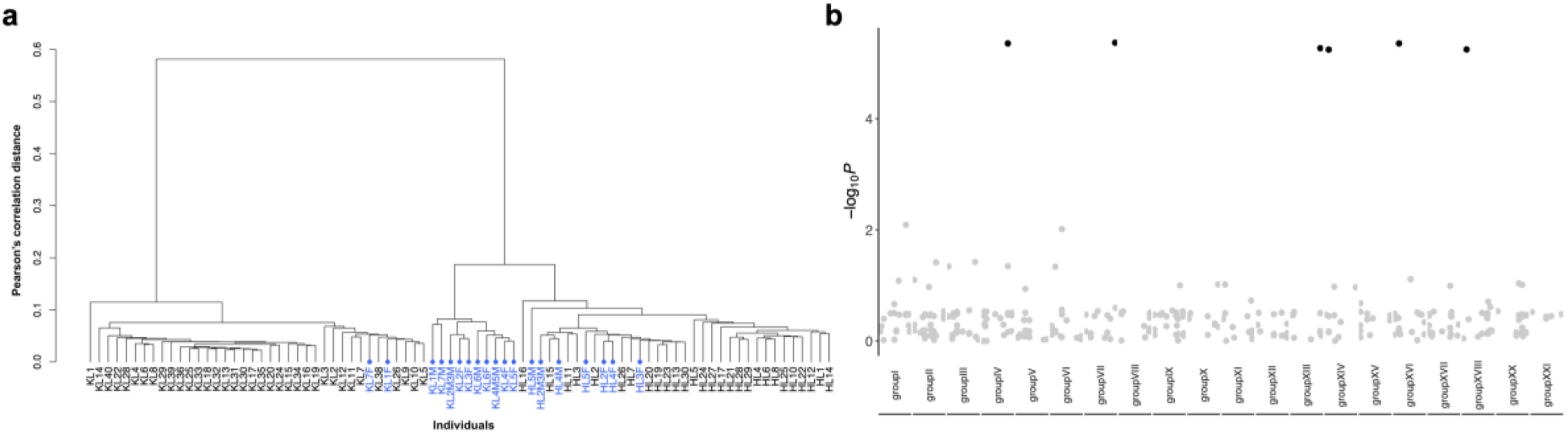
(a) Dendrogram of methylation levels for all fish in F1 and F2 generations. The y-axis is the Pearson’s correlation distance after hierarchical clustering of the percent methylation levels of the 52,729 intergenerationally stable CpG sites. F1 fish are shown in blue, and F2 fish are shown in black. (b) Manhattan plot showing the -log*P* of correlations between each single nucleotide polymorphism (SNPs) (columns) and the 52,729 intergenerationally stable CpG sites when filtering SNPs using a 10% missing data cut-off. Black points are statistically significant SNPs (*Q* < 0.05) after adjusting for multiple testing using the Benjamini-Hochberg’s false discovery rate method. SNPs (n = 43) from unassembled scaffolds are without significant hits, and thus are not shown here.

We found no significant enrichment of the stable sites in any of the genomic contexts when compared to the null distribution of all filtered sites (promoters: *G* = 4.53 × 10^-4^, df = 1, *P* = 0.983; exons: *G* = 4.94 × 10^-5^, df = 1, *P* = 0.994; introns: *G* = 3.49 × 10^-5^, df = 1, *P* = 0.995; intergenic regions: *G* = 1.01 × 10^-4^, *P* = 0.992; Fig. 5a). Among the stable CpG sites, we found 6,462, 28,005 and 18,262 CpG sites that were constitutively hypermethylated, constitutively hypomethylated, and variable, respectively. When analysing the genomic context of CpGs from these three categories, we found a significantly biased genomic distribution, with constitutively hypermethylated sites enriched within exons (*G* = 29.3, df = 1, *P* = 6.33 × 10^-8^; Fig. 5b), constitutively hypomethylated sites enriched within promoters (*G* = 17.8, df = 1, *P* = 2.42 × 10^-5^; Fig. 5c), and variable sites enriched within introns (*G* = 9.36, df = 1, *P* = 2.22 × 10^-3^; Fig. 5d) when compared to the null distribution of all filtered CpGs. We found that 94.8% (845 out of 891) of the DMCs between ecotypes were also identified as stable sites, a percentage not significantly different than the percentage of stable sites among all filtered sites (*G* = 0.171, df = 1, *P* = 0.679), suggesting that most sites involved in methylation divergence between ecotypes can be stable across generations, and could therefore be plausibly associated with adaptation to different habitats.

**Fig. 5.**
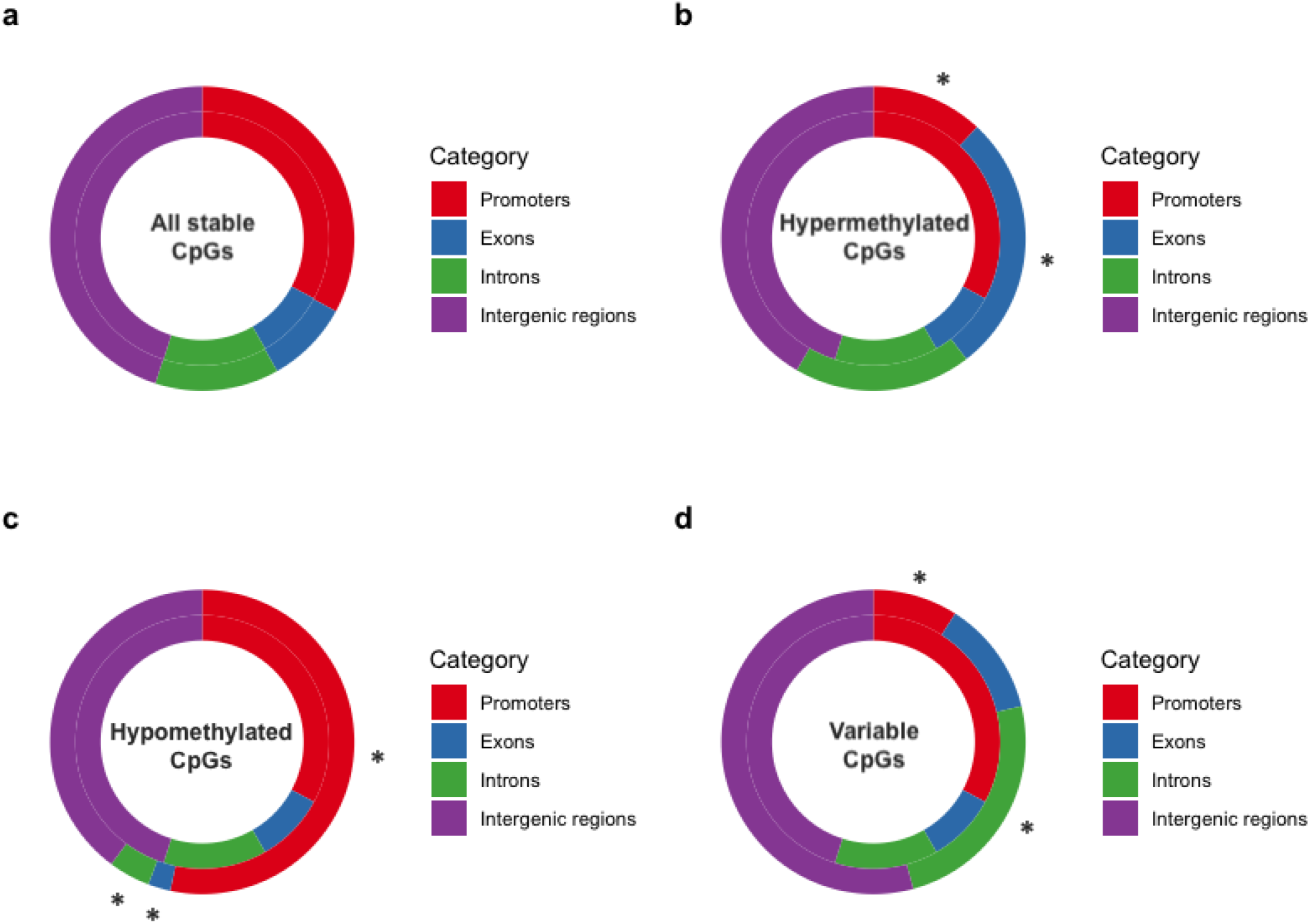
The proportion of genomic features (promoters, exons, introns or intergenic regions) in intergenerationally stable CpGs compared with null distribution of all filtered CpGs in (a) all intergenerationally stable sites, (b) constitutively hypermethylated sites, (c) constitutively hypomethylated sites, and (d) variable sites. Outer rings describe the locations of CpGs in each category; inner rings describe the features of null distribution of all filtered CpGs. Asterisks denote significant differences between the features of CpGs in each category vs. the features of null distribution of CpGs across the genome using a *G* test at *P* < 0.01.

To calculate the genetic heritability of stable sites, we first identified 92,983 SNPs, and then filtered this dataset down to 350 SNPs that 1) had less than 10% missing data across all F1 and F2 individuals, 2) were not located on the sex chromosome, and 3) had low to no LD with each other. Six of these SNPs showed highly significant associations with the methylation values of F1 and F2 individuals (*Q* < 0.05), and 16,514 out of the 52,729 intergenerationally stable CpG sites had h^2^ > 0 (Fig. 4b). We also retained 3,007 and 4,203 SNPs after filtering the SNPs by 30% and 50% missing data, respectively, with 22 and 28 of these SNPs showing highly significant associations with the methylation values of F1 and F2 individuals, and 16,498 and 21,055 intergenerationally stable CpG sites having h^2^ > 0 (Fig. S5). Finally, the kinship matrix estimated the narrow sense heritability for CpG methylation levels at (on average) 24%, 32% and 35% using the 350, 3,007 and 4,203 post-filtering SNPs, respectively.

### Identification of meQTLs associated with methylation divergence between marine and freshwater ecotypes

When analyzing meQTLs within each hybrid line, we identified 968 and 531 significant SNP-CpG pairs in HL_F2 and KL_F2 fish, respectively, corresponding to 335 unique SNPs and 75 unique CpG sites in HL_F2, and 201 unique SNPs and 72 unique CpG sites in KL_F2. We found that 85.0% (HL_F2: 823 of 968) and 94.4% (KL_F2: 501 out of 531) of the SNP-CpG pairs were also identified as significant SNP-CpG pairs when using R/qtl, and the *P*-value distributions showed no evidence of test statistic inflation (Fig. S6). A two-dimensional plot of meQTLs indicates that each SNP could regulate multiple CpG sites located across the genome (*trans*-meQTLs indicated as grey dots show large scatter around the diagonal line of *cis*-meQTLs, indicated as black dots; Fig. S7). We found meQTLs displayed significantly more *trans-* than *cis*-meQTLs (HL_F2: 965 *trans-* vs. 3 *cis*-meQTLs, *G* = 1.30 × 10^3^, df = 1, *P* < 2.20 × 10^-16^; KL_F2: 522 *trans-* vs. 4 *cis*-meQTLs, *G* = 689, df = 1, *P* < 2.20 × 10^-16^), with no significant GO term enrichment for genes annotated with unique SNPs within significant meQTLs in either line. There was relatively low LD between SNPs and CpGs with significant *cis* associations (mean r^2^ = 0.287). We also found no overlap between CpGs associated with meQTLs and the DMCs identified between ecotypes in HL_F2, likely due to the small number of CpG sites associated with significant meQTLs. Alternatively, this could be due to the small amount (ranging from 24% to 35%) of methylation variation explained by genetic variation, suggesting that a significant proportion of CpGs are likely to be autonomous from genetic variation, and thus not detectable in a meQTL analysis. We found two CpGs associated with nine *trans-*meQTLs that overlapped with DMCs in KL_F2, with all nine SNP-CpG pairs verified by R/qtl. Although the two CpGs did not localise within any genes, they were in close genomic proximity (~15 kb) to zinc finger E-box binding homeobox 1b *(zeb1b,* Ensembl Gene ID ENSGACG00000001002) and centrosomal protein 76 *(cep76,* Ensembl Gene ID ENSGACG00000003686). The nine *trans*-meQTLs were annotated with four genes (Table S3).

To assess the co-occurrence of meQTLs and eQTLs, we compared locations of meQTLs identified in our study to eQTL hotspots in Ishikawa *et al.* (2017). We found that an overall of 9.14% (HL_F2: 24 out of 335; KL_F2: 25 out of 201) of the unique SNPs overlapped with eQTL locations across HL_F2 and KL_F2 samples. This proportion does not suggest an excess of meQTL-eQTL overlap relative to null expectations that are built from all input SNPs for meQTL analyses (*G* = 0.245, df = 1, *P* = 0.621). Finally, to investigate whether meQTLs might be associated with divergent selection in marine vs. freshwater habitats, we examined whether the unique SNPs within meQTLs overlapped with genomic regions of high differentiation between the two stickleback ecotypes. We found a total of six (four in HL_F2 and two in KL_F2) unique SNPs within high differentiation regions, corresponding to 14 (11 and 3 in HL_F2 and KL_F2, respectively) *trans-*acting SNP-CpG pairs. The effect sizes (beta) of meQTLs ranged from 0.105 to 0.406 (median = 0.161). The six SNPs were annotated with four genes (Table S4), which encode proteins likely to be relevant to marine-freshwater divergence in stickleback (e.g., sensing changes in osmoregulatory environment; see below).

## Discussion

The role of DNA methylation in fundamental ecological and evolutionary processes has received increased attention in recent years (Metzger & Schulte 2016; Verhoeven *et al.* 2016; Hu & Barrett 2017). However, the extent to which variation in DNA methylation is stably transmitted across generations, and the prevalence of *cis*-and/or *trans*-acting genetic variants in contributing to methylome evolution remain poorly understood, particularly in natural animal populations. We used a quantitative, single-base-resolution technique (RRBS) to measure DNA methylation from fin tissue across two generations in threespine stickleback sampled from two distinct environment types. A large majority (99.6%) of CpG sites were identified as being intergenerationally stable, as indicated by consistent methylation levels across F1 and F2 generations in two hybrid lines. As a consequence, of the subset of CpG sites that also showed significant divergence between marine and freshwater ecotypes in the grandparental generation, a large majority (94.8%) could be classified as being stable across generations. Epigenetic variation was associated with genetic variation to some extent, with a narrow sense heritability ranging from 24% to 35%. These values are consistent with recent epigenome-wide association studies that have found that genetic variation can explain an average of 7-34% of methylation variation in animals (McRae *et al.* 2014; Orozco *et al.* 2015; Taudt *et al.* 2016; Carja *et al.* 2017). We found distinct patterns of genomic context between three categories of stable CpG sites: constitutively hypomethylated and hypermethylated sites were predominantly located within promoters and exons, respectively, whereas variable sites were enriched within introns. We also identified meQTLs in marine-freshwater F2 hybrid lines, with some meQTLs overlapping with genomic regions of high differentiation between marine and freshwater ecotypes in stickleback. Finally, we identified the genetic architecture underlying two DMCs between ecotypes that were also shown to have intergenerational stability in their methylation levels. Overall, our study provides the first investigation of the genetic basis of stable epigenetic variation in stickleback and identifies methylation differences that could be associated with local adaptation in marine and freshwater ecotypes.

### Methylation divergence between ecotypes

We found a similar number of differential methylation sites (891 DMCs) between marine and freshwater ecotypes of threespine stickleback to a recent study (737 DMCs in Smith *et al.* 2015). While we did not find any significantly enriched GO terms, some of these DMCs were annotated with genes that are likely to contribute to adaptation to marine and freshwater environments. For example, we found DMCs overlapped with genes related to osmoregulation (ion channel activity: *trpc1, RYR3, gria3b, kcnq3),* metabolic process (lipid and fatty acid metabolism: *elovl6l*, *scap*; glucose and carbohydrate metabolism: *g6pd*), immune response (hemopoiesis: *kalrna;* myeloid cell and neutrophil differentiation: *satb1b;* erythrocyte maturation: *klf3*), and catalytic activity *(alpl, phlpp1, sdr39u1).* Because the osmotic environments, parasite communities, and migratory life cycles of marine and freshwater ecotypes differ (Smith *et al.* 2015; Huang *et al.* 2016; Artemov *et al.* 2017; Ishikawa *et al.* 2017), the differential methylation of these genes suggests that the methylome could be associated with ecologically important phenotypic differentiation between ecotypes.

### The genetic basis and functions of intergenerationally stable epigenetic variation

To understand how methylation divergence between ecotypes might be involved in the process of local adaptation, we next explored the stability of epigenetic variation across generations. While the approach of using experimental crosses to explore stable epigenetic variation and its underlying genetic basis has been widely applied in plant studies (e.g., Johannes *et al.* 2009; Roux *et al.* 2011; Li *et al.* 2014), very few studies have used this type of experimental design in non-model animal populations (but see Nätt *et al.* 2012; Weyrich *et al.* 2016; Weyrich *et al.* 2018). Examination of the genetic basis of methylation sites is valuable for exploring the mechanisms that facilitate animal responses to novel environments, and for predicting the likelihood that populations will be able to evolve in response to environmental change (O’Dea *et al.* 2016). We identified 52,729 CpG sites that were not differentially methylated between F1 and F2 generations in both HL and KL hybrid lines (99.6% of all CpG sites). Similar to Heckwolf *et al.* (2020), this suggests that the majority of our analysed CpG sites have stable levels of methylation across generations. These CpG sites are not necessarily heritable; it is possible that methylation levels are induced to similar levels across generations due to exposure to a similar environment. This pattern may also be due to non-global DNA methylation reprogramming during embryogenesis in fish, which can provide greater opportunity for transmitting DNA methylation from parents to the offspring (Schmitz *et al.* 2011; Skvortsova *et al.* 2018).

When assessing the contribution of the intergenerationally stable CpG sites to evolutionary processes, we found significant enrichment of constitutively hypomethylated CpG sites within promoters, and significant enrichment of constitutively hypermethylated CpG sites within gene bodies, suggesting that stable DNA methylation may directly regulate gene expression and facilitate alternative splicing, and thus contribute to genomic evolution by providing access to alternative promoter sites and increasing the number of transcriptional opportunities and phenotypes (Roberts & Gavery 2012). Consequently, mutations impacting intergenerationally stable methylation could accelerate the exploration of phenotypic space, and therefore allow populations to adapt to the changing environments more efficiently (Klironomos *et al.* 2013). Furthermore, different categories of stable sites showed distinct features of genomic context, with hypermethylated sites enriched within exons and constitutively hypomethylated sites enriched within promoters. The distinct distribution patterns of hyper- and hypomethylated CpG sites are consistent with whole genome assessments of methylation in other fish species and in model animals and plants (Feng *et al.* 2010; Zemach *et al.* 2010; Long *et al.* 2013; Shao *et al.* 2014), and suggest a conserved role for constitutive hyper- and hypomethylation in a wide range of species.

We found a large proportion (~95%) of DMCs between marine and freshwater ecotypes in the grandparental generation also showed intergenerational stability in methylation levels, suggesting that the genes associated with these DMCs could play a role in facilitating adaptation to different environments. Theoretical work has suggested that environmentally responsive epigenetic changes that can be transmitted to the next generation can be beneficial when the effects of epigenetic variation increase both parental and offspring fitness with low cost (Herman *et al.* 2014). As the functions of several DMC-associated genes identified here are relevant to responses to changes in aquatic environments such as salinity, parasites, and diet, our findings provide evidence for a possible adaptative mechanism in threespine stickleback whereby advantageous epigenetic changes that have been triggered by environmental stimulus are transmitted across generations.

There is substantial interest in biomedical and agricultural fields to understand the contribution of genetic variation to population epigenomic variation, with a number of recent genetic studies having quantified the heritable basis of population epigenomic variation in model animals (Taudt *et al.* 2016). When applying a stringent missing data cut-off (10%), we found a reasonably high average heritability of 24% for methylation levels across F1 and F2 generations.

When applying more relaxed missing data cut-offs (30% and 50%), we find heritability estimates of 32% and 35%. In addition, we found that 31% to 40% of stable CpG sites had a measurable genetic component (narrow-sense heritability h^2^ > 0), a percentage similar to previous findings in model species (Taudt *et al.* 2016). Together, our results suggest a plurality of mechanisms are likely contributing to stable levels of methylation variation across generations, including genetic control, epimutation, and exposure to past or current environmental factors.

### Contribution of meQTL to methylation divergence between ecotypes

We characterised meQTLs in the F2 generation of two marine-freshwater hybrid lines, and detected two CpGs associated with significant meQTLs that overlapped with DMCs between marine and freshwater ecotypes, both in the KL_F2 line. The two CpGs were close to *zeb1b* and *cep76,* which are key genes involved in developmental processes. *Zeb1b* has been hypothesized to be a regulator of interleukin 2, which is associated with differences in parasite load of stickleback inhabiting marine and freshwater environments, and consequently affects their immune responses (Scharsack *et al.* 2016; Verta & Jones 2019). *Cep76* is an important paralog of *CC2D2A,* which is a gene associated with development of the primary cilia, and is relevant to morphological differences between Pacific lamprey populations with distinct migratory behaviours (Hess *et al.* 2014). Because the morphology between marine and freshwater stickleback ecotypes differs significantly (Jones *et al.* 2012), the overlap between meQTL-associated CpGs and ecotype-DMCs suggests that these loci may be under divergent selection in marine versus freshwater habitats.

Although methylation variation between marine and freshwater ecotypes can be caused by both *cis*- and *trans-*regulatory changes, we found only *trans*-meQTLs within genomic regions of high differentiation between ecotypes. This is interesting because we expected to detect a bias towards *cis*-meQTLs due to the close genomic proximity of the SNPs and CpGs from the same RRBS fragments. Moreover, a number of recent studies have shown greater contribution of *cis*-regulatory than *trans-*regulatory genetic variants in gene expression divergence in the gill, brain and liver tissues of stickleback (Ishikawa *et al.* 2017; Pritchard *et al.* 2017; Verta & Jones 2019). However, predominantly *trans-*regulatory changes in gene expression have also been found in the tooth plate of stickleback and *Drosophila* (McManus *et al.* 2010; Osada *et al.* 2017; Hart *et al.* 2018). These contrasting results have been attributed to a number of factors such as inter-vs. intra-specific comparison and tissue heterogeneity, where *tran*s-regulatory effects dominate in intraspecific comparisons and in more heterogeneous tissue (Hart *et al.* 2018). Our findings fit with these explanations in that we conducted an intraspecific comparison using heterogenous caudal fin tissue that consists of epidermis, osteoblasts, dermal fibroblasts, and vascular endothelium (Tu & Johnson 2011).

Although selection may initially favour master regulator genes that regulate distant genes through *trans*-acting mechanisms during rapid adaptation, it has been suggested that different evolutionary scenarios and selective contexts may alternatively favour *trans-* and *cis*-acting mechanisms during intraspecific adaptive divergence (Cooper *et al.* 2003; Lemos *et al.* 2008; Stern & Orgogozo 2009; Hart *et al.* 2018). In the case of marine stickleback invading freshwater environments, local adaptation must often occur in the presence of ongoing gene flow (Nelson & Cresko 2018). This scenario may initially favour *trans*-acting mechanisms that are less susceptible to being eroded through recombination. However, as the population reaches later stages of adaptation to the local environment, the advantage of responses mediated by *trans-* regulatory genes may shift to favour *cis*-regulatory mechanisms, where co-evolved mutations are more closely linked to each other and the genes they regulate (Verta & Jones 2019).

Our functional analysis identified multiple genes associated with meQTLs that are located in genomic regions that have been shown to have significant differentiation between marine and freshwater populations, and could therefore be relevant for local adaptation (Hohenlohe *et al.* 2010; Jones *et al.* 2012; Terekhanova *et al.* 2014) (Table S4). For example, we found genes annotated with osmosis and electrolyte transport *(KCNB2’),* and skeletal and fibroblast growth *(Slco5a1a),* which have also been found in previous stickleback studies investigating differential gene expression or methylation in gill or fillet tissue between the marine and freshwater ecotypes (Smith *et al.* 2015; Artemov *et al.* 2017). These results suggest that this collection of genes might be important for facilitating adaptation to these divergent environments in stickleback, although the direction of causality between DNA methylation variation and gene expression remain elusive. Interestingly, it has been shown that genomic regions that are not significantly differentiated between ecotypes can still play an important role in adaptation to novel aquatic environments in stickleback (DeFaveri *et al.* 2011; Leinonen *et al.* 2012; Ellis *et al.* 2015; Erickson *et al.* 2016; Ferchaud & Hansen 2016). Our findings suggest that the genetic architecture underlying methylation divergence and physiological adaptation to different aquatic environments in stickleback is complicated and could include SNPs from genomic regions that experience either neutral or selective processes.

### Limitations

Our study has a number of caveats that should be noted. First, an intrinsic problem of *in vivo* studies using next-generation sequencing techniques such as RRBS is the heterogeneity of analysed tissues. Fin tissues consist of many different cell types including epidermis, osteoblasts, dermal fibroblasts, and vascular endothelium (Tu & Johnson 2011). Therefore, various proportions of different cell types could introduce biases in measures of methylation levels (Kratochwil & Meyer 2015). In addition, we only used fin tissues from a single developmental stage of sticklebacks, whereas methylation and gene expression patterns are known to be development-related and tissue-specific (Wang *et al.* 2009; Feil & Fraga 2012), and thus, overlap between the locations of meQTLs identified in this study and eQTLs in Ishikawa *et al.* (2017) should be interpreted with caution. Further studies extending our work to a broader range of tissues and developmental stages will be helpful for a more comprehensive characterisation of methylation variation, and its role in gene regulation and development.

Second, the reduced representation genome sequencing method used here can only cover a small proportion of all possible methylation patterns in these populations. Thus, we are inevitably missing a large number of stable CpG sites and SNPs located outside of the regions of the genome represented here. In addition, because the accuracy of SNP calls from bisulfite sequencing data can be affected by the conversion rate of unmethylated cytosines (Barturen *et al.* 2014), the SNPs identified in our study could be different than those that would be obtained using a sequencing method that produces independent SNP data (e.g., restriction-site associated DNA sequencing, RAD-seq; Baird *et al.* 2008). Furthermore, while we mainly focused methylation patterns in gene bodies, other regulatory elements such as enhancers and transposons, although less well annotated in stickleback, are also important drivers of regulatory and phenotypic evolution (Wittkopp & Kalay 2011) and thus warrant further research.

Third, the number of individuals and families used in our study is limited. Thus, including additional samples from more families would provide additional information on family-level variation, as well as more loci associated with methylation variation that would increase the power of our heritability and meQTL analyses. In addition, we generated marine-freshwater F1 families by crossing marine females and freshwater males, and recent studies in zebrafish have suggested that epigenetic patterns at early developmental stages can often reprogram to reflect the paternal state (Jiang *et al.* 2013; Potok *et al.* 2013). Whether such reprogramming is common to teleosts remains unclear (Skvortsova *et al.* 2018), but additional reciprocal crosses using marine males and freshwater females, as well as pure crosses within marine or freshwater populations, would allow a more comprehensive understanding of parental effects on epigenetic inheritance (Laporte *et al.* 2019).

Fourth, while we found *cis* associations between SNPs and CpGs, this does not necessarily indicate that the CpGs are under genetic control. Although average LD between SNPs and CpG sites with significant *cis* associations was relatively low, it is still possible that patterns are driven in part by linkage disequilibrium between epigenetic variation at a locus and its proximal SNP (Taudt *et al.* 2016; Heckwolf *et al.* 2020). It is also possible that the CpGs are autonomous from genetic control and contribute to heritable variation that is shaped by natural selection, and thus will be indistinguishable from genetic variation in a standard heritability analysis (Johannes *et al.* 2008; Helanterä & Uller 2010; Tal *et al.* 2010). Thus, the results of our stable methylation and meQTL analyses should be interpreted with consideration of these alternatives.

Finally, although we have corrected for the possibility of falsely interpreting C-to-T and G-to-A SNPs as epigenetic variation by excluding them from methylation estimates, it is possible that some SNPs were miscalled. Thus, our results provide a necessarily coarse map of the genetic architecture underlying stable methylation and methylation divergence between marine and freshwater stickleback populations. A wider investigation of regulatory elements in combination with genome-wide sequencing of chromatin modifications (e.g., chromatin immunoprecipitation followed by sequencing (ChIP-seq); Park 2009; Furey 2012) and whole-genome resequencing (e.g., Le Luyer *et al.* 2017) would provide a more comprehensive and precise understanding of the relationship between genetics and DNA methylation, and the role that epigenetic responses may play in facilitating evolutionary change.

## Conclusions

Here, we provide the first insights into the genetic architecture of DNA methylation in threespine stickleback. Our genome-wide methylation data reveals that the vast majority of CpG sites have stable methylation levels across generations, including the sites that show significant divergence in methylation levels between marine and freshwater ecotypes. Some of these sites show evidence of genetic control, while others are likely to be autonomous from genetic variation. We also explored the genomic distribution of methylation in marine-freshwater hybrid populations and found meQTLs that overlap with previously identified genomic regions of high differentiation between marine and freshwater populations. In addition, our data demonstrates different contributions of *cis*- and *trans*-meQTLs to methylome divergence in stickleback. Our study adds to the few studies using non-model, outbred vertebrates to test for the genetic basis of intergenerationally stable methylation and methylation divergence between ecotypes. Our results suggest that methylation could play an important role in facilitating phenotypic plasticity over the short-term, as well as population persistence and adaptation over longer evolutionary time scales.

## Acknowledgements

The authors would like to thank Antoine Paccard and Caroline LeBlond for their assistance in sample preparation, and Paul Gugger for his guidance in performing association analysis. This work was supported by China Scholarship Council Fellowship 201406350023 and Start-up research funding from Fudan University to JH, Queen Elizabeth II Scholarships to SJSW and TNB, Alberta Innovates to TNB, a National Sciences and Engineering Research Council (NSERC) RT735278 to SMR, a NSERC Discovery Grant 418249-2012 to HAJ, and a NSERC Discovery Grant 429955-2013 and Canada Research Chair to RDHB. We would like to acknowledge the Bamfield Marine Sciences Centre for resources in the collection of fishes and during the writing of this manuscript.

## Author Contributions

JH and RDHB conceived the study. SJSW, TB, HAJ, and SMR sampled, crossed and reared fish, and collected tissue. JH generated and analysed sequencing data. JH wrote the manuscript with input from SMR, and RDHB. The authors have no conflicts of interest.

## Notes

### Competing Interest Statement

The authors have declared no competing interest.

