## supplemental material for "Heritability of DNA methylation in threespine stickleback (*Gasterosteus aculeatus*)"

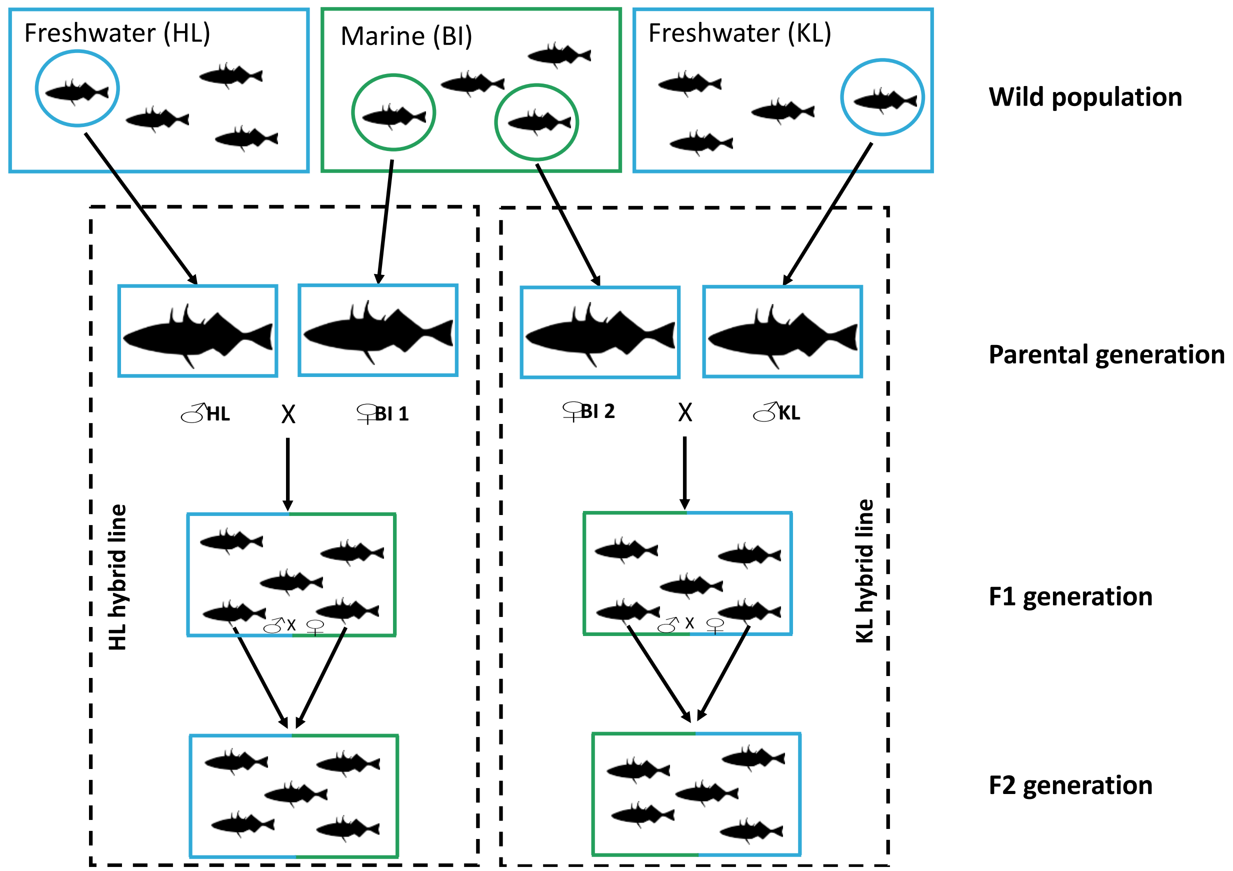


**Fig. S1** Overview of the experimental design for this study. Adult threespine stickleback from one marine (BI) and two freshwater (HL and KL) locations were collected and transplanted to our aquatic facility. F1 hybrid lines were produced by randomly selecting and crossing one male fish from HL or KL and one female fish from BI. In total, one F1 family of HL hybrids was produced, and three F1 families of KL hybrids were produced (two additional F1 families of KL hybrids are not shown here). To generate F2 families, one male and one female sibling within an F1 family were randomly selected and crossed from each hybrid line (HL or KL), resulting in one F2 family of HL hybrids and one F2 family of KL hybrids.

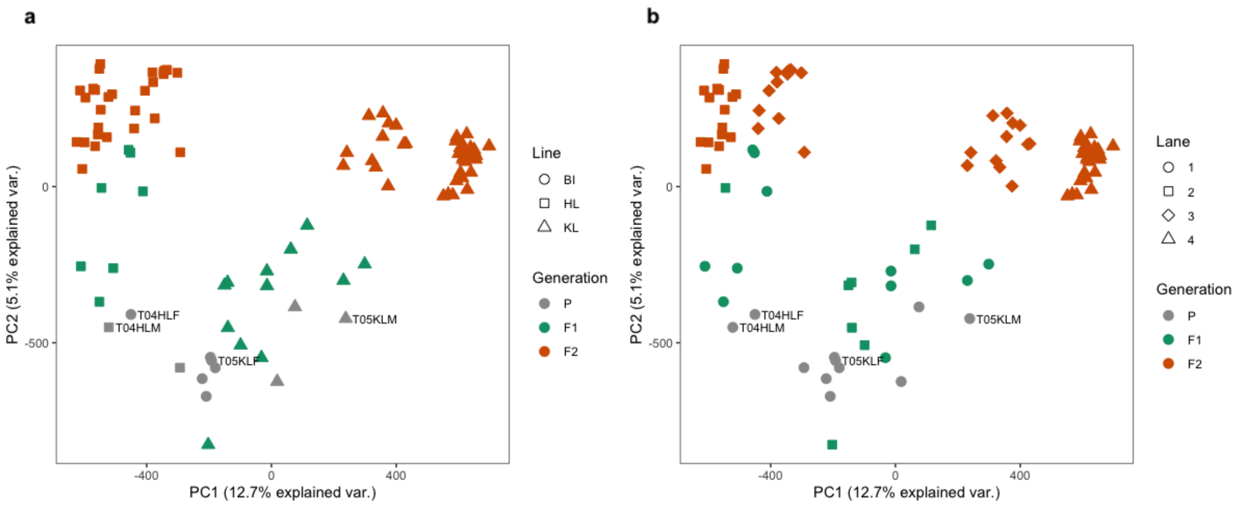


**Fig. S2** Principal component analysis (PCA) of DNA methylation profiles based on all CpG sites after filtering in all individuals from parental, F1 and F2 generations. PCA axes explain 12.7% (PC1) and 5.1% (PC2) of the total variation. a) Sampling site of parental fish in generation P, parental sire of fish in the F1 generation, and grandparental sire of fish from the F2 generation. Lane: b) Sequencing lane. The parents of the F1 families used to produce the F2 generation are labelled (see Table S1).

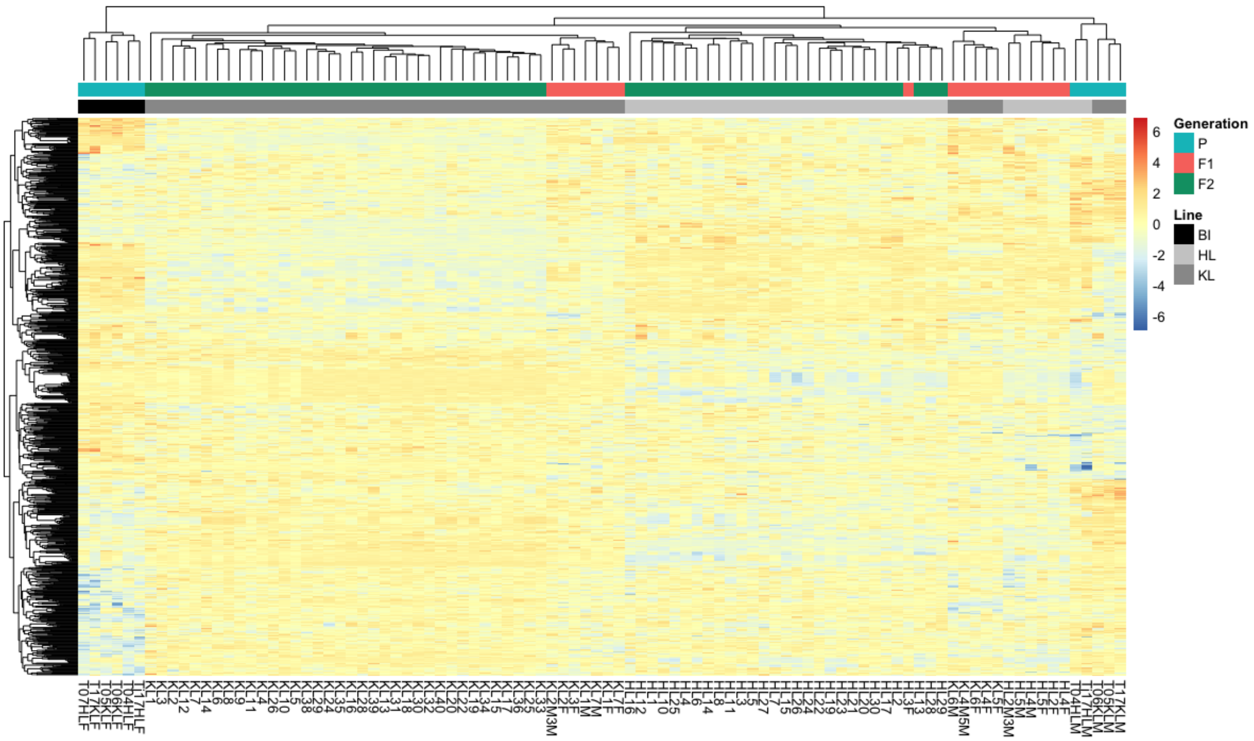


**Fig. S3** Heatmap of methylation levels of the 891 CpGs that are identified as DMCs between marine and freshwater ecotypes from the parental generation across parental, F1, and F2 generations. Each column represents a colour-coded individual: blue for parental fish, red for F1 fish, green for F2 fish; black for marine fish from BI, light grey for freshwater fish from HL or hybrid HL lines, and dark grey for freshwater fish from KL or KL hybrid lines. Each row represents one of the CpGs, which are clustered based on the similarities of the methylation patterns between individuals. Darker red indicates greater methylation in an individual for that DMC. Darker blue indicates lesser methylation in an individual for that CpG. Individual dendrogram positions are based on overall methylation patterns across the 891 DMCs.

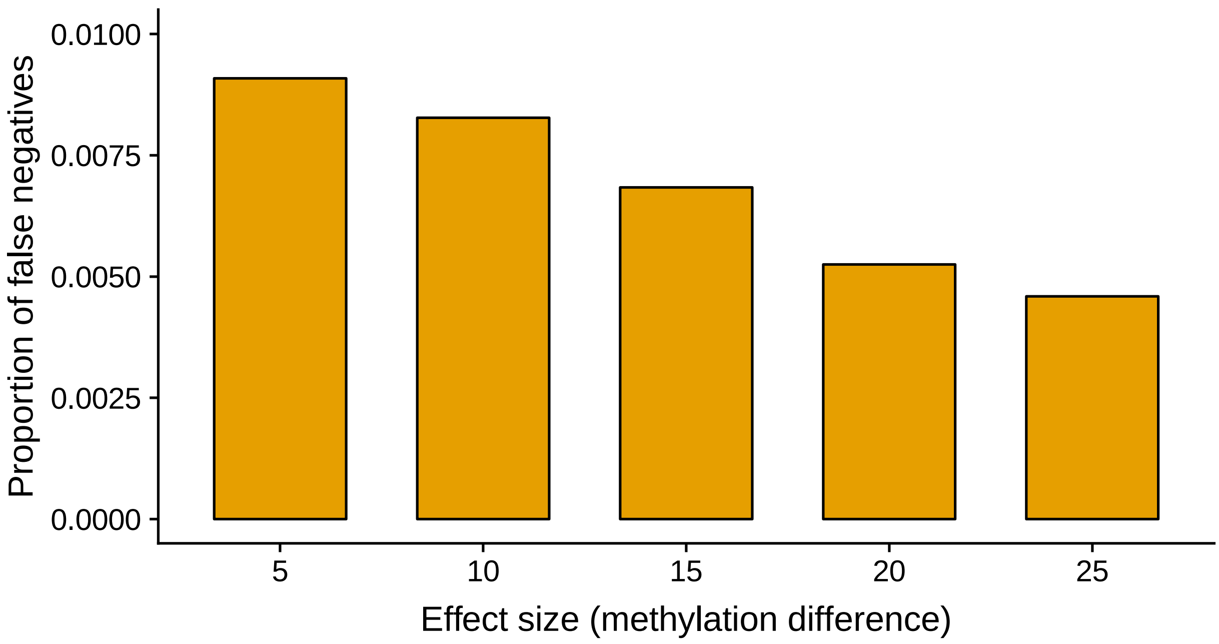


**Fig. S4** Proportion of falsely identified non-DMC (false negatives) among all CpG sites in simulated data. Effect size indicates methylation differences between two groups of samples (F1s and F2s). Replicates in one group had methylation changes in 1% of CpGs sites by 5%, 10%, 15%, 20% and 25%, respectively.

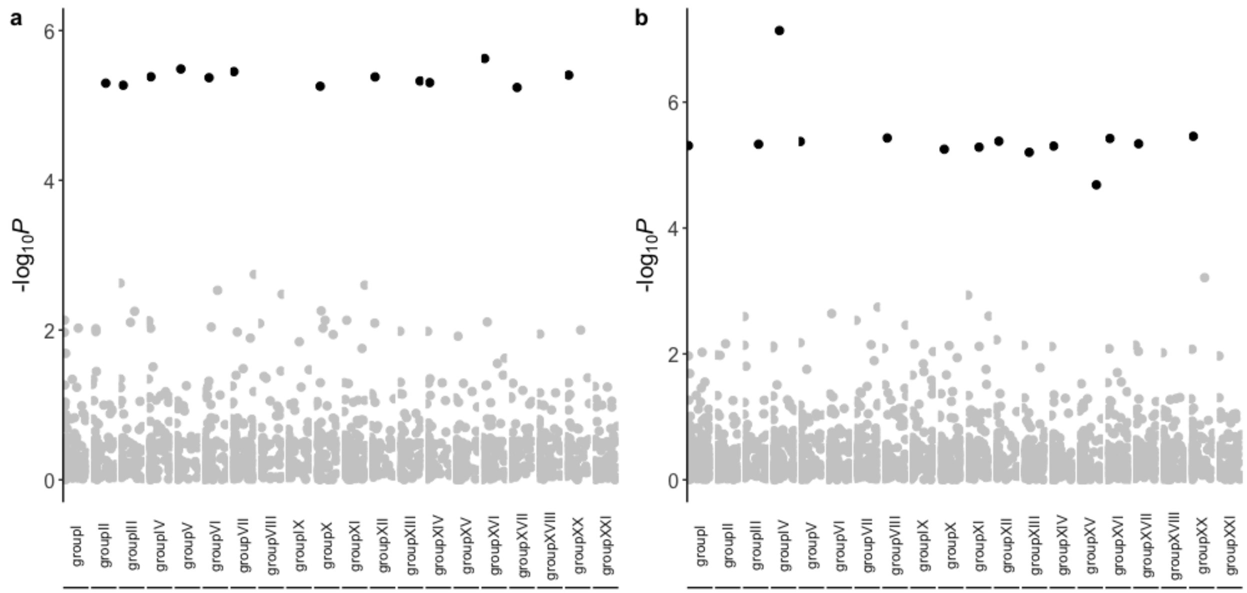


**Fig. S5** Manhattan plot showing the -log*P* of correlations between each single nucleotide polymorphism (SNPs) (columns) and the 52,729 intergenerationally stable CpG sites when filtering SNPs using (a) 30% and (b) 50% missing data cut-off. Black points are statistically significant SNPs ($Q < 0.05$) after adjusting for multiple testing using the Benjamini-Hochberg's false discovery rate method. SNPs (n = 9 for 30% cut-off, n = 14 for 50% cut-off) from unassembled scaffolds with significant associations with the methylation values of F1 and F2 individuals are not shown here.

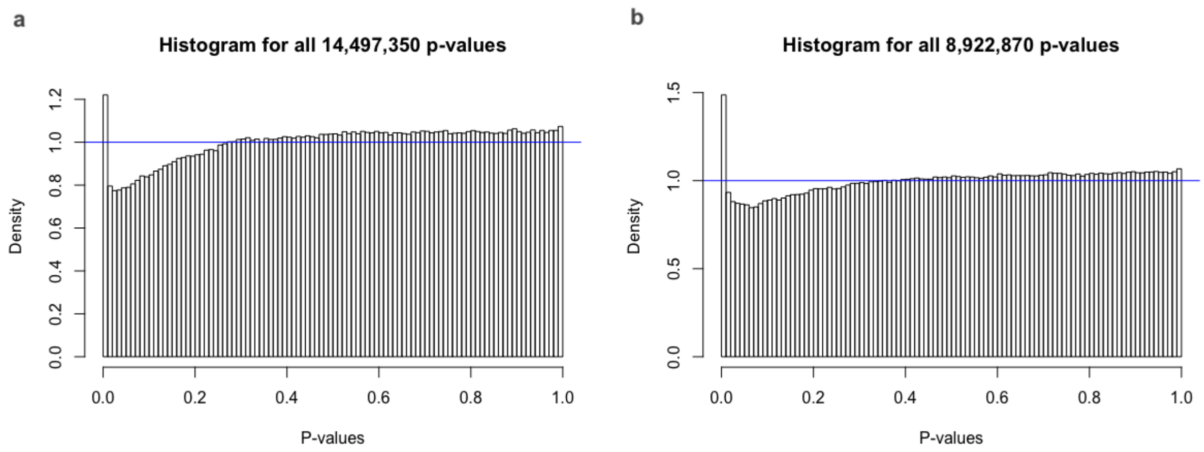


**Fig. S6** *P*-value distribution of meQTL analysis in (a) HL_F2 fish and (b) KL_F2 fish.

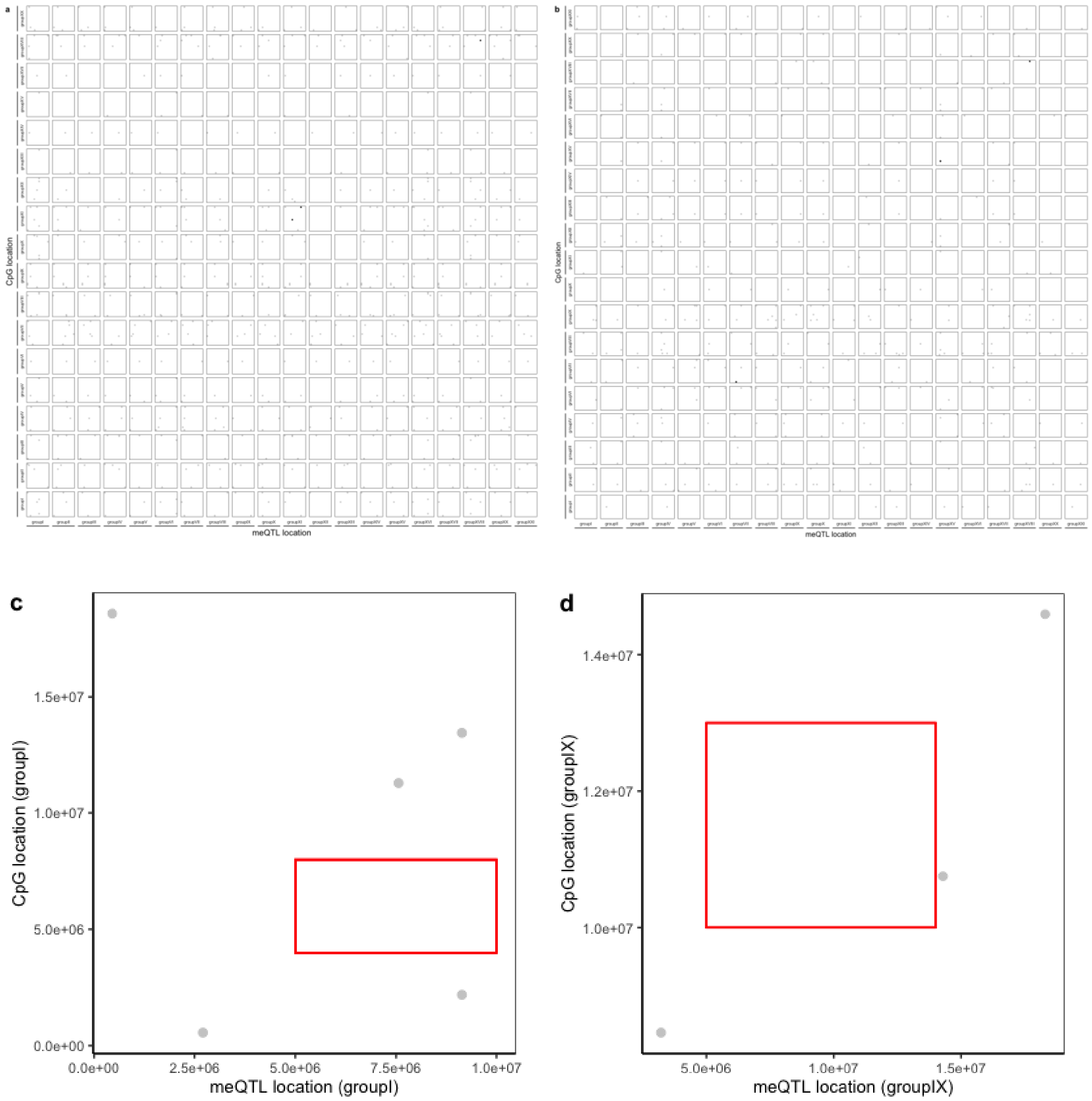


**Fig. S7** Significant *cis*- and *trans*-methylation quantitative trait loci (meQTL) in (a) HL_F2 and (b) KL_F2. Each dot represents a single meQTL on chromosome. Black dots along the diagonal line show *cis*-meQTLs, whereas grey dots indicate *trans*-meQTLs. A zoomed in plot showing significant *cis*- and *trans*-meQTL (c) in linkage group I in HL_F2 and (d) in linkage group IX in KL_F2. Blocks of white space indicate non-significant associations between SNPs and CpGs. Red squares in panel (c-d) show examples of blocks associated with non-significant meQTLs.

**Table S1** Samples used in this study

| Sample ID | Sex[1] | Sequencing lane | Generation | Wild habitat | Parental/Grandparental sire[2] | Family |
|---|---|---|---|---|---|---|
| **T05KLF** | F | 1 | Parental | Marine | N/A | N/A |
| **T05KLM** | M | 1 | Parental | Freshwater | N/A | N/A |
| T07HLF | F | 1 | Parental | Marine | N/A | N/A |
| **T04HLF** | F | 1 | Parental | Marine | N/A | N/A |
| **T04HLM** | M | 1 | Parental | Freshwater | N/A | N/A |
| T06KLF | F | 1 | Parental | Marine | N/A | N/A |
| T06KLM | M | 1 | Parental | Freshwater | N/A | N/A |
| T17KLF | F | 1 | Parental | Marine | N/A | N/A |
| T17KLM | M | 1 | Parental | Freshwater | N/A | N/A |
| Tj17HLF | F | 1 | Parental | Marine | N/A | N/A |
| Tj17HLM | M | 1 | Parental | Freshwater | N/A | N/A |
| **KL1F** | F | 1 | F1 | N/A | KL | KL_F1_1 |
| **KL1M** | M | 1 | F1 | N/A | KL | KL_F1_1 |
| HL4M | M | 1 | F1 | N/A | HL | HL_F1_1 |
| HL4F | F | 1 | F1 | N/A | HL | HL_F1_1 |
| **HL2M3M** | M | 1 | F1 | N/A | HL | HL_F1_1 |
| **HL2F** | F | 1 | F1 | N/A | HL | HL_F1_1 |
| HL3F | F | 1 | F1 | N/A | HL | HL_F1_1 |
| KL2M3M | M | 1 | F1 | N/A | KL | KL_F1_2 |
| KL2F | F | 1 | F1 | N/A | KL | KL_F1_2 |
| KL3F | F | 1 | F1 | N/A | KL | KL_F1_2 |
| HL5M | M | 1 | F1 | N/A | HL | HL_F1_1 |
| HL5F | F | 2 | F1 | N/A | HL | HL_F1_1 |
| KL7F | F | 2 | F1 | N/A | KL | KL_F1_1 |
| KL7M | M | 2 | F1 | N/A | KL | KL_F1_1 |
| KL6F | F | 2 | F1 | N/A | KL | KL_F1_3 |
| KL6M | M | 2 | F1 | N/A | KL | KL_F1_3 |

| | | | | | | |
|---|---|---|---|---|---|---|
| KL4M5M | M | 2 | F1 | N/A | KL | KL_F1_3 |
| KL5F | F | 2 | F1 | N/A | KL | KL_F1_3 |
| KL4F | F | 2 | F1 | N/A | KL | KL_F1_3 |
| HL1 | F | 2 | F2 | N/A | HL | HL_F2_1 |
| HL2 | F | 2 | F2 | N/A | HL | HL_F2_1 |
| HL3 | M | 2 | F2 | N/A | HL | HL_F2_1 |
| HL4 | M | 2 | F2 | N/A | HL | HL_F2_1 |
| HL5 | F | 2 | F2 | N/A | HL | HL_F2_1 |
| HL6 | N/A | 2 | F2 | N/A | HL | HL_F2_1 |
| HL7 | F | 2 | F2 | N/A | HL | HL_F2_1 |
| HL8 | F | 2 | F2 | N/A | HL | HL_F2_1 |
| HL10 | F | 2 | F2 | N/A | HL | HL_F2_1 |
| HL11 | M | 2 | F2 | N/A | HL | HL_F2_1 |
| HL12 | F | 2 | F2 | N/A | HL | HL_F2_1 |
| HL13 | N/A | 2 | F2 | N/A | HL | HL_F2_1 |
| HL14 | F | 2 | F2 | N/A | HL | HL_F2_1 |
| HL15 | M | 2 | F2 | N/A | HL | HL_F2_1 |
| HL16 | M | 2 | F2 | N/A | HL | HL_F2_1 |
| HL17 | F | 2 | F2 | N/A | HL | HL_F2_1 |
| HL19 | M | 3 | F2 | N/A | HL | HL_F2_1 |
| HL20 | M | 3 | F2 | N/A | HL | HL_F2_1 |
| HL21 | F | 3 | F2 | N/A | HL | HL_F2_1 |
| HL22 | M | 3 | F2 | N/A | HL | HL_F2_1 |
| HL23 | F | 3 | F2 | N/A | HL | HL_F2_1 |
| HL24 | F | 3 | F2 | N/A | HL | HL_F2_1 |
| HL25 | M | 3 | F2 | N/A | HL | HL_F2_1 |
| HL26 | M | 3 | F2 | N/A | HL | HL_F2_1 |
| HL27 | F | 3 | F2 | N/A | HL | HL_F2_1 |
| HL28 | F | 3 | F2 | N/A | HL | HL_F2_1 |

| | | | | | | |
|---|---|---|---|---|---|---|
| HL29 | N/A | 3 | F2 | N/A | HL | HL_F2_1 |
| HL30 | N/A | 3 | F2 | N/A | HL | HL_F2_1 |
| KL1 | F | 3 | F2 | N/A | KL | KL_F2_1 |
| KL2 | N/A | 3 | F2 | N/A | KL | KL_F2_1 |
| KL3 | M | 3 | F2 | N/A | KL | KL_F2_1 |
| KL4 | F | 3 | F2 | N/A | KL | KL_F2_1 |
| KL5 | M | 3 | F2 | N/A | KL | KL_F2_1 |
| KL6 | M | 3 | F2 | N/A | KL | KL_F2_1 |
| KL7 | F | 3 | F2 | N/A | KL | KL_F2_1 |
| KL8 | M | 3 | F2 | N/A | KL | KL_F2_1 |
| KL9 | N/A | 3 | F2 | N/A | KL | KL_F2_1 |
| KL10 | M | 3 | F2 | N/A | KL | KL_F2_1 |
| KL11 | M | 3 | F2 | N/A | KL | KL_F2_1 |
| KL12 | M | 3 | F2 | N/A | KL | KL_F2_1 |
| KL13 | M | 4 | F2 | N/A | KL | KL_F2_1 |
| KL14 | M | 4 | F2 | N/A | KL | KL_F2_1 |
| KL15 | F | 4 | F2 | N/A | KL | KL_F2_1 |
| KL16 | F | 4 | F2 | N/A | KL | KL_F2_1 |
| KL17 | M | 4 | F2 | N/A | KL | KL_F2_1 |
| KL18 | F | 4 | F2 | N/A | KL | KL_F2_1 |
| KL19 | M | 4 | F2 | N/A | KL | KL_F2_1 |
| KL20 | M | 4 | F2 | N/A | KL | KL_F2_1 |
| KL22 | M | 4 | F2 | N/A | KL | KL_F2_1 |
| KL24 | M | 4 | F2 | N/A | KL | KL_F2_1 |
| KL25 | M | 4 | F2 | N/A | KL | KL_F2_1 |
| KL26 | F | 4 | F2 | N/A | KL | KL_F2_1 |
| KL28 | F | 4 | F2 | N/A | KL | KL_F2_1 |
| KL29 | M | 4 | F2 | N/A | KL | KL_F2_1 |
| KL30 | M | 4 | F2 | N/A | KL | KL_F2_1 |

| | | | | | | | |
|---|---|---|---|---|---|---|---|
| KL31 | M | 4 | F2 | N/A | KL | | KL_F2_1 |
| KL32 | M | 4 | F2 | N/A | KL | | KL_F2_1 |
| KL33 | F | 4 | F2 | N/A | KL | | KL_F2_1 |
| KL34 | M | 4 | F2 | N/A | KL | | KL_F2_1 |
| KL35 | M | 4 | F2 | N/A | KL | | KL_F2_1 |
| KL36 | M | 4 | F2 | N/A | KL | | KL_F2_1 |
| KL38 | F | 4 | F2 | N/A | KL | | KL_F2_1 |
| KL39 | M | 4 | F2 | N/A | KL | | KL_F2_1 |
| KL40 | F | 4 | F2 | N/A | KL | | KL_F2_1 |

[1]Sex: 'F' is female, 'M' is male

[2]Parental/Grandparental sire: parental sire denotes sire of F1 generation, grandparental sire denotes sire of F2 generation.

**Bold blue** indicates the parents and grandparents of KL fish in F2 generation, and **bold red** indicates the parents and grandparents of HL fish in F2 generation

**Table S2** CpG sites with top 10 lowest and highest loadings on PC1 when analysing all samples

| Chromosome/Scaffold | Location | PC1 loading |
|---|---|---|
| scaffold_139 | 11217 | -0.070907 |
| groupXX | 16586212 | -0.0653622 |
| groupIX | 7428278 | -0.05766 |
| groupVIII | 18090060 | -0.0571543 |
| groupVII | 26539349 | -0.0552824 |
| groupXVIII | 3405322 | -0.0549021 |
| groupXX | 2311484 | -0.0534399 |
| groupI | 4749376 | -0.0504257 |
| scaffold_129 | 64293 | -0.0500742 |
| scaffold_129 | 64292 | -0.0491858 |
| scaffold_184 | 10898 | 0.04698332 |
| scaffold_121 | 292637 | 0.04745949 |
| groupXV | 2267 | 0.04778125 |
| groupXVII | 11605505 | 0.05384838 |
| groupXX | 16586225 | 0.05693498 |
| groupIV | 29366481 | 0.05758826 |
| groupXII | 261435 | 0.05797566 |
| groupXVII | 12294190 | 0.06453433 |
| groupVIII | 14481109 | 0.06473624 |
| groupIX | 17906031 | 0.07059687 |

**Table S3** List of genes annotated with the nine meQTLs underlying the DMC between marine and freshwater ecotypes

| Ensembl Gene ID | Symbol | GO ID | GO Term | Description |
|---|---|---|---|---|
| ENSGACG00000001561 | bsnb | GO:0007416 | synapse assembly | bassoon (presynaptic cytomatrix protein) b [Source:ZFIN;Acc:ZDB-GENE-120628-1] |
| ENSGACG00000001561 | bsnb | GO:0045202 | synapse | bassoon (presynaptic cytomatrix protein) b [Source:ZFIN;Acc:ZDB-GENE-120628-1] |
| ENSGACG00000001561 | bsnb | GO:0046872 | metal ion binding | bassoon (presynaptic cytomatrix protein) b [Source:ZFIN;Acc:ZDB-GENE-120628-1] |
| ENSGACG00000001561 | bsnb | GO:0048786 | presynaptic active zone | bassoon (presynaptic cytomatrix protein) b [Source:ZFIN;Acc:ZDB-GENE-120628-1] |
| ENSGACG00000001561 | bsnb | GO:0042995 | cell projection | bassoon (presynaptic cytomatrix protein) b [Source:ZFIN;Acc:ZDB-GENE-120628-1] |
| ENSGACG00000002303 | spegb | GO:0005886 | plasma membrane | striated muscle enriched protein kinase b [Source:ZFIN;Acc:ZDB-GENE-081104-517] |
| ENSGACG00000002303 | spegb | GO:0016020 | membrane | striated muscle enriched protein kinase b [Source:ZFIN;Acc:ZDB-GENE-081104-517] |
| ENSGACG00000002303 | spegb | GO:0016021 | integral component of membrane | striated muscle enriched protein kinase b [Source:ZFIN;Acc:ZDB-GENE-081104-517] |
| ENSGACG00000002303 | spegb | GO:0004672 | protein kinase activity | striated muscle enriched protein kinase b [Source:ZFIN;Acc:ZDB-GENE-081104-517] |
| ENSGACG00000002303 | spegb | GO:0004674 | protein serine/threonine kinase activity | striated muscle enriched protein kinase b [Source:ZFIN;Acc:ZDB-GENE-081104-517] |
| ENSGACG00000002303 | spegb | GO:0005524 | ATP binding | striated muscle enriched protein kinase b [Source:ZFIN;Acc:ZDB-GENE-081104-517] |
| ENSGACG00000002303 | spegb | GO:0006468 | protein phosphorylation | striated muscle enriched protein kinase b [Source:ZFIN;Acc:ZDB-GENE-081104-517] |
| ENSGACG00000002303 | spegb | GO:0000166 | nucleotide binding | striated muscle enriched protein kinase b [Source:ZFIN;Acc:ZDB-GENE-081104-517] |
| ENSGACG00000002303 | spegb | GO:0005515 | protein binding | striated muscle enriched protein kinase b [Source:ZFIN;Acc:ZDB-GENE-081104-517] |
| ENSGACG00000000768 | NA | GO:0005576 | extracellular region | NA |
| ENSGACG00000000768 | NA | GO:0016787 | hydrolase activity | NA |
| ENSGACG00000000768 | NA | GO:0006508 | proteolysis | NA |

| | | | | |
|---|---|---|---|---|
| ENSGACG00000000768 | NA | GO:0004252 | serine-type endopeptidase activity | NA |
| ENSGACG00000000768 | NA | GO:0008236 | serine-type peptidase activity | NA |
| ENSGACG00000000768 | NA | GO:0005520 | insulin-like growth factor binding | NA |
| ENSGACG00000000768 | NA | GO:0005515 | protein binding | NA |

**Table S4** List of genes annotated with meQTLs in HL_F2 or KL_F2 fish that are located within genomic regions of high differentiation between marine and freshwater ecotypes in Hohenlohe *et al.* (2010), Jones *et al.* (2012), and Terekhanova *et al.* (2014)

| Ensembl Gene ID | Symbol | Description |
|---|---|---|
| HL_F2 | | |
| ENSGACG00000002793 | *KCNB2* | potassium voltage-gated channel subfamily B member 2 |
| ENSGACG00000002831 | *Slco5a1a* | solute carrier organic anion transporter family member 5A1 |
| KL_F2 | | |
| ENSGACG00000018951 | NA | NA |
| ENSGACG00000019342 | NA | NA |
